# A Synthetic Platform for Antibody Junctional Diversification Beyond Natural Constraints

**DOI:** 10.64898/2026.05.21.726110

**Authors:** Wang Liu, Jun Yang, Yuhang Zhang, Xiaoli Xue

**Author notes:** Corresponding authors: Xiaoli Xue; Yuhang Zhang.

## Abstract

Antibody junctional diversity in jawed vertebrates arose from RAG transposon domestication, generating CDR3-focused V(D)J recombination with lymphocyte specificity—a constrained system limiting engineered antibody diversity. To overcome these limitations, we developed ARESEC (**A**ntibody **R**eprogram and **E**xpression **S**ystem in **E**ngineered **C**ells), a synthetic platform that enables programmable V(D)J recombination in non-lymphoid HEK293T cells. Through engineered recombination signal sequences (RSS) and optimized RAG1/2 expression, ARESEC enables RSS-guided DNA recombination across all three complementarity-determining regions (CDR1/2/3). Co-expression of terminal deoxynucleotidyl transferase (TdT) enhanced junctional sequence diversity by 41.5-86.3% across 3 CDRs. The platform enables native IgG production by coupling mammalian surface display with FACS-based functional screening. Using clinical-grade antibodies (Nivolumab and Durvalumab) as high-affinity starting scaffolds, we achieved further efficient affinity maturation through focused CDR diversification (library complexity >10^3^). From these diversification libraries, we identified optimized variants including the anti-PD-L1 Dur1 with 2.01-fold enhanced binding, demonstrating the system’s capacity for rapid antibody optimization from minimal diversity sampling. Collectively, ARESEC establishes a synthetic paradigm that transcends natural V(D)J constraints, generating multi-CDR diversification in non-lymphoid cells to enable rapid discovery of affinity-matured antibodies, effectively bridging immune evolution with modern antibody engineering demands.

## 1. Introduction

Adaptive immunity in jawed vertebrates originated with the emergence of RAG-mediated V(D)J recombination^[1]^. This somatic DNA rearrangement enables the combinatorial assembly of antigen receptor genes, and the imprecise joining of gene segments further generates extensive junctional diversity^[2]^. Together, these evolutionary innovations provide the vast repertoire necessary for adaptive immune recognition. This mechanism is mediated by the RAG recombinase, which evolved from an ancestral RAG-like transposon comprising RAG1L and RAG2L. Following invasion into an immunoglobulin superfamily (*IgSF*) gene and suppression of transposition activity, this genetic element gave rise to the RAG1/RAG2 recombinase system. Whole-genome duplication and subsequent tandem duplications of *Ig* gene segments then contributed to the expansion and diversification of antibody and T-cell receptor (TCR) loci in jawed vertebrates (∼500 MYA)^[3]^. The primary repertoire is further diversified through TdT-mediated N-nucleotide addition and Artemis-dependent P-nucleotide formation^[4, 5]^. While lymphocytes utilize this system to produce antibodies or TCRs, non-immune somatic cells lack the inherent ability to perform V(D)J recombination and generate functional antigen receptors (**Figure 1**). ^[6]^

**Figure 1.**
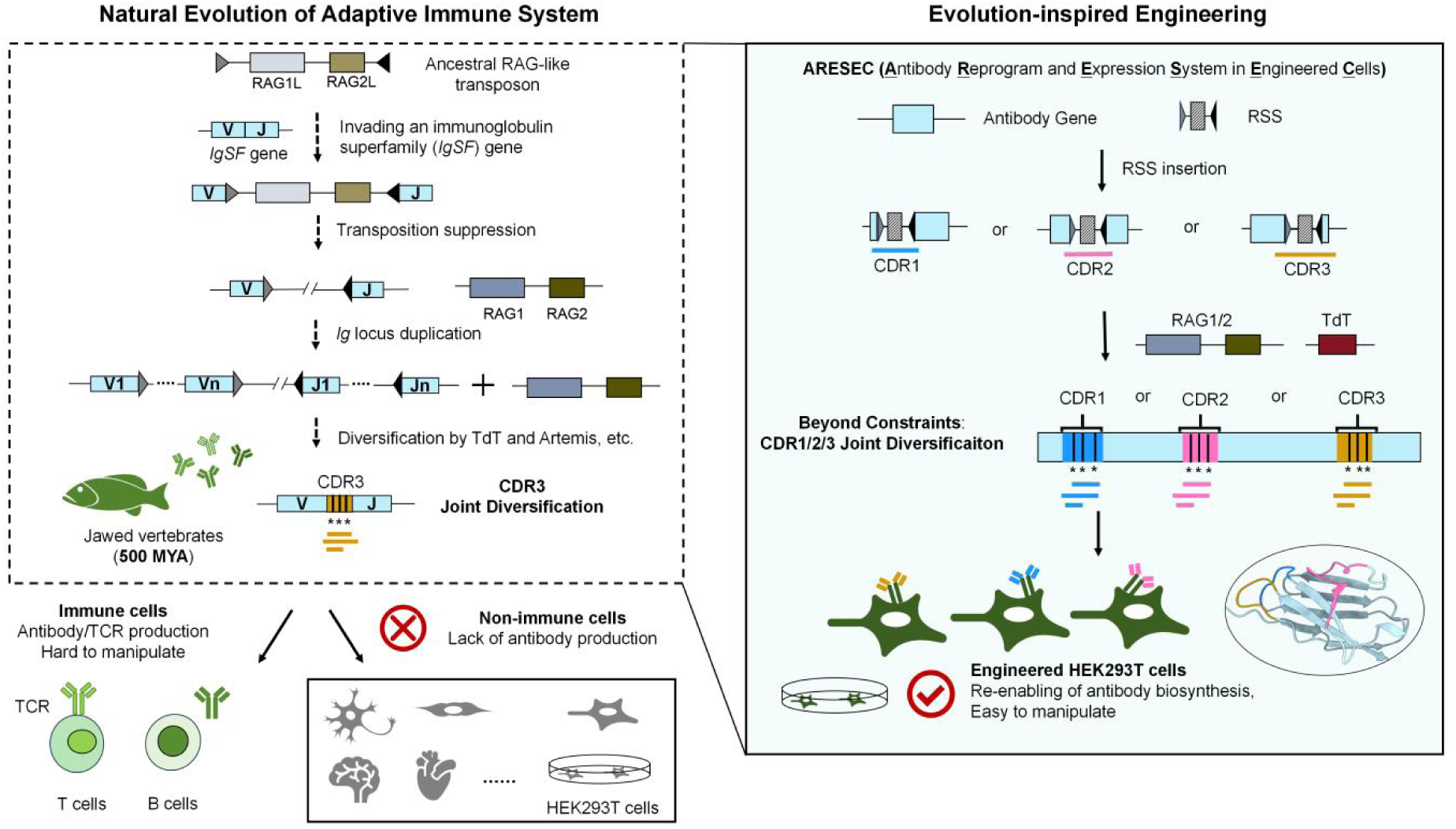
Synthetic expansion of antibody diversity beyond evolutionary constraints. Left panel: Natural evolution of the adaptive immune system. The adaptive immune system in jawed vertebrates relies on V(D)J recombination, a lymphocyte-specific DNA rearrangement process, to generate diverse antigen receptor repertoires. This mechanism is catalyzed by the RAG recombinase, which originated from an ancestral RAG-like transposon harboring both RAG1L and RAG2L modules. Following its insertion into an immunoglobulin superfamily (*IgSF*) gene and subsequent suppression of transposition activity, this element evolved into the canonical RAG1/RAG2 recombinase system. Whole-genome duplication and subsequent tandem duplications of *Ig* gene segments drove the expansion and diversification of antibody and T-cell receptor (TCR) loci. These gene segments undergo diversification via junctional processing mediated by TdT and Artemis, further expanding the antibody repertoire. Co-evolution of the RAG system enabled the catalysis of V(D)J recombination specifically in lymphocytes, thereby establishing adaptive immunity. Notably, the junctional diversity generated through V(D)J recombination is structurally confined to the CDR3 region of antigen receptors. This evolutionary design constrains the primary structural diversification of the antigen-binding site to the CDR3 loop. Following the divergence of immune and non-immune lineages, the RAG-mediated diversification machinery was silenced in somatic cells, rendering them incapable of *de novo* antibody production. Right panel: Reconstruction of antibody diversification in engineered somatic cells using the ARESEC platform. The ARESEC (Antibody Reprogram and Expression System in Engineered Cells) platform introduces recombinant signal sequences (RSS) into CDR1, CDR2, and CDR3 regions of antibody gene segments. Co-expression of RAG1/2 and TdT in engineered HEK293T cells enables site-specific recombination and nucleotide addition, thereby extending junctional diversification from its evolutionarily restricted CDR3 compartment to include CDR1 and CDR2. This expansion of diversification capacity across all three CDR loops enables the *de novo* biosynthesis of highly diverse antibody repertoires in a tractable non-immune cellular chassis. This synthetic approach recapitulates an evolutionarily latent capacity, effectively overcoming the natural constraint of CDR3-limited diversity and enabling programmable antibody biosynthesis in somatic cells.

In B lymphocytes, immunoglobulin diversity arises via: (i) the combinatorial V(D)J rearrangement^[7]^; (ii) P/N nucleotide addition during V(D)J recombination^[8]^; (iii) combinatorial pairing of heavy and light chains^[9]^; and (iv) SHM-mediated sequence refinement^[10]^. SHM in B cells introduces C to T/G to A transitions preferentially in antibody CDRs (indel frequency **<**5%)^[11]^, driven by AID deamination of single-stranded DNA. In contrast, T cell receptors (TCRs) diversify solely through V(D)J recombination and junctional modifications, lacking SHM due to AID restriction to B cells^[12]^.

V(D)J recombination, mediated by the RAG1/RAG2 recombinase^[13]^, is a fundamental mechanism shared by both B and T lymphocytes to generate the vast diversity of antigen receptors essential for adaptive immunity^[1]^. The RAG1/RAG2 recombinase orchestrates V(D)J recombination by recognizing conserved RSS motifs (heptamer/nonamer with 12/23-bp spacers) to initiate DNA cleavage^[14] [15] [16]^, generating combinatorial diversity and junctional flexibility at CDR3^[17]^. Notably, The C-terminal domain of RAG2 independently regulates locus-specific *Igκ* demethylation in pre-B cells-a critical epigenetic precondition for recombination. The C-terminal truncated mutant of RAG2 could increase recombinase activity *in vivo* ^[18] [19]^. RAG-mediated double-strand breaks are resolved by NHEJ, while CDR3 diversity is further amplified by TdT-dependent N-additions, Artemis-mediated P-nucleotide processing, and nucleolytic trimming^[20] [21]^. This stochastic cascade produces extreme CDR3 sequence variability, establishing its role as the primary antigen-contact determinant^[22] [23] [24]^.

Although the natural immune system can generate a broad repertoire of antibodies, the evolutionarily shaped sequence space does not fully meet the optimization requirements of therapeutic antibodies in terms of affinity, specificity, and pharmacokinetics^[25] [26] [27]^; therefore, further engineering and functional enhancement of existing antibodies may be a key strategy to improve their efficacy and expand their range of applications^[28]^.

Current antibody engineering platforms face fundamental trade-offs between throughput and biological fidelity. Cell-free systems (e.g., ribosome display) enable ultrahigh-throughput screening but are limited to the use of structurally incomplete single-chain variable fragments (scFv) as antibody formats and lack the capacity for post-translational modifications^[29] [30]^; microbial systems (e.g., yeast/phage display) permit rapid affinity maturation yet cannot replicate native CDR3 length variation (5**–**30 aa) or glycosylation patterns^[31] [32] [33]^; while mammalian systems (e.g., hybridomas) preserve physiological heavy-light chain pairing and glycosylation, they remain constrained by host repertoire limitations^[34] [35] [36] [37]^. These persistent gaps, particularly in replicating natural CDR plasticity and cognate VH-VL pairing, underscore the need for next-generation synthetic biology solutions using human cellular platforms ^[38] [39]^.

The synthetic biology paradigm provides transformative capabilities for biological engineering. Advances in DNA synthesis, assembly, and AI-guided design^[40] [41] [42]^ now enable whole genome construction, accelerated through automated biofoundries implementing design-build-test cycles^[43]^. This framework has been successfully adapted for antibody engineering. The CellectAb platform ^[44]^ integrates CRISPR-mediated CDR1 targeting with Jurkat cells’ endogenous error-prone repair and ER-retention selection, generating native-like antigens against antigens (e.g., CD3), representing a significant advancement in synthetic immunology. Similarly, the PnP-HDM system ^[45]^ employs homology-directed mutagenesis (15**–**35% HDR efficiency) in hybridoma cells for targeted antibody diversification through CRISPR/Cas9 and degenerate codon templates, enabling mammalian display screening for full-length IgG optimization through targeted diversity generation. While these immune cell-based systems represent breakthroughs in synthetic immunology, their reliance on T cells/hybridomas introduces inherent limitations-low genetic tractability, costly culturing, and poor scalability-that motivate the development of engineered non-immune cell platforms for programmable antibody discovery and production ^[46] [47]^.

Building upon the evolutionary foundations of adaptive immunity, we engineered ARESEC (Antibody Reprogramming and Expression System in Engineered Cells)-a synthetic biology platform that reconstitutes and enhances V(D)J recombination in the human HEK293T cell line (**Figure 1**). This system not only recapitulates the core RAG-mediated recombination machinery, but also expands its diversification capacity beyond natural constraints through engineered RSS sequences and TdT co-expression. By establishing this programmable antibody biosynthesis platform in a tractable somatic cell model, we demonstrate efficient generation and screening of affinity-matured variants, and successful applications in therapeutic antibody optimization.

## 2. Result

### 2.1. Development and optimization of ARESEC in HEK293T Cells

We developed ARESEC (Antibody Reprogramming and Expression System in Engineered Cells) to enable antibody gene diversification in non-lymphoid HEK293T cells through systematic integration of three key evolution-inspired components (**Figure 1**) : (i) programmable RSS site integration following the conserved 12/23 rule; (ii) regulated expression of engineered RAG recombinase that maintains core enzymatic functions while overcoming lymphocyte-specific constraints; and (iii) TdT-mediated junctional diversification extending beyond natural N-addition to enhance combinatorial diversity. This synthetic system achieves unprecedented expansion of V(D)J-mediated diversification, generating hypervariable regions not only in CDR3 but also in CDR1 and CDR2. Integrated with mammalian surface display technology, ARESEC supports both native IgG production and high-throughput functional screening of affinity-matured variants.

To validate the feasibility of engineered V(D)J recombination, we selected the Hepatitis B surface antigen (HBSAg) antibody light chain (AB159729.1) as a model system. The expression construct was designed with flanking recombination signal sequences RSS and an intervening SV40 poly(A) signal to suppress transcription of unrearranged antibody genes. Co-transfection of HEK293T cells with the VL-RSS-polyA-RSS-JL-CL-FLAG and RAG1/RAG2 expression plasmids yielded full-length light chain expression (25 kDa) by western blot (**Figure 2A**), confirming successful V-J recombination. To quantify recombination efficiency, we developed an EGFP-based reporter system, with flow cytometric analysis showing 1.19% EGFP-positive cells, whereas control transfections with mutated RSS (12/23RSSmut) or catalytically inactive RAG1 (RAG^mut^) exhibited only background fluorescence (Figure S1). These results establish that both proper RSS recognition and functional RAG enzymatic activity are essential for efficient recombination in non-lymphoid cells.

**Figure 2.**
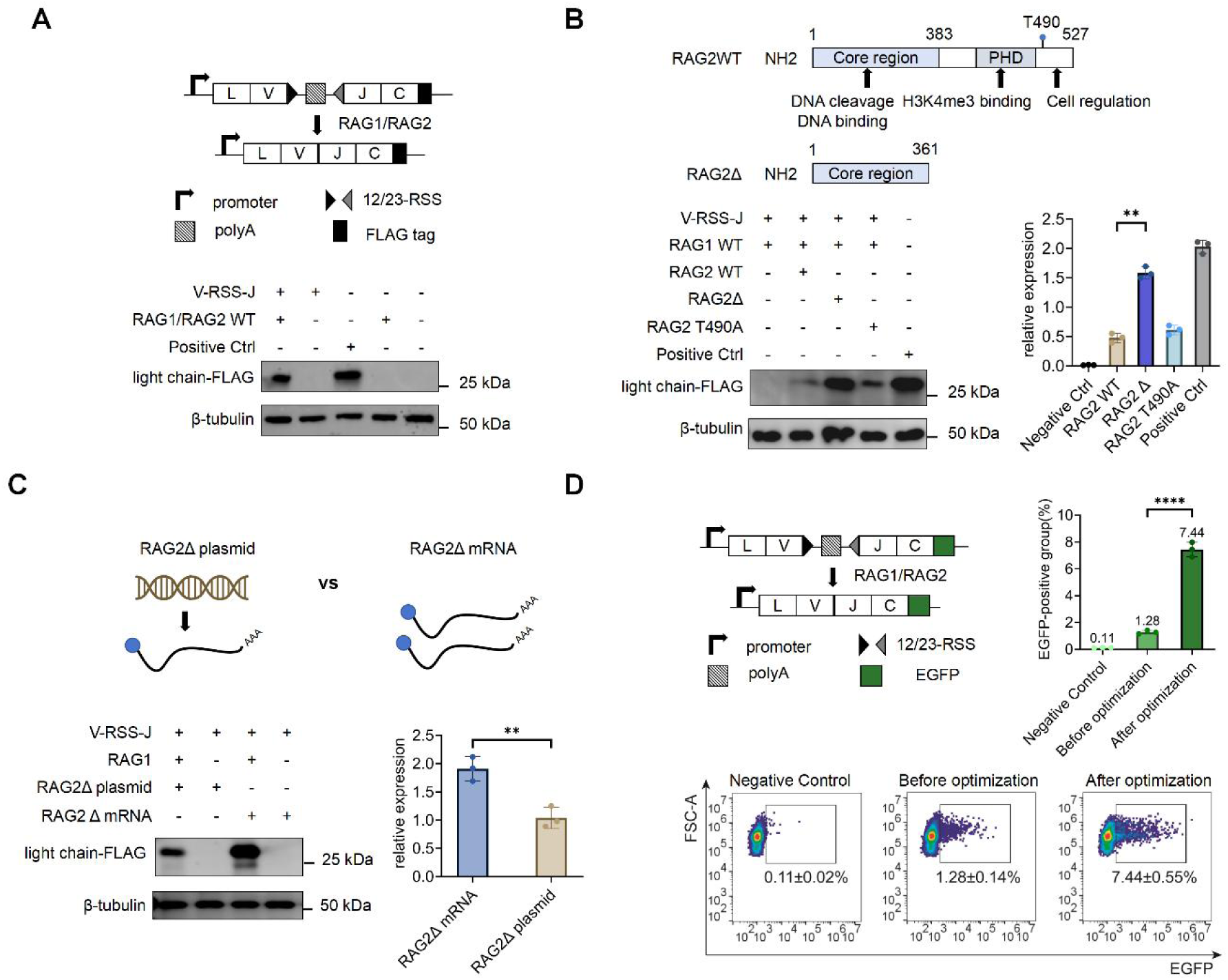
Optimization of RAG1/2-mediated antibody gene rearrangement in HEK293T cells. (A) Western blot analysis of antibody light chain expression following gene rearrangement mediated by RAG1/2. Positive control: plasmid containing correctly-expressing V-J sequence. (B) Schematic representation of the wild-type and truncated RAG2, and Western blot analysis of rearrangement products according to different RAG2 derivatives. Negative control: no RAG2; Positive control: plasmid containing correctly-expressing V-J sequence. (C) Western blot analysis of rearrangement products after transfection with either plasmid DNA or mRNA encoding RAG2Δ. (D) Flow cytometric analysis of recombination efficiency. HEK293T cells were transfected with the reporter plasmid and RAG1/2. Left: negative control, no RAG1/2; middle: before optimization; right: after optimization.

RAG2 expression in the innate immune system is spatiotemporally regulated via its plant homeodomain (PHD) and CDK2-mediated T490 phosphorylation^[3]^ ^[19]^ ^[48]^, confining V(D)J recombination to G1 phase for genomic stability ^[49] [50]^. To enhance V(D)J recombination efficiency in HEK293T cells, we systematically validated RAG2 mutants. The C-terminal truncated mutant (RAG2Δ, residues 1-361) significantly improved recombination efficiency (**Figure 2B**), while the full-length phosphorylation site mutant (RAG2 T490A) showed only mild enhancement. Further optimization revealed that replacing plasmid transfection with mRNA delivery significantly enhanced light chain expression **(Figure 2C**), leading to a 5.8-fold improvement in recombination efficiency (from 1.28% to 7.44% EGFP-positive cells; **Figure 2D**). Co-expression of HMGB1 with RAG1/2, though proposed to enhance RAG-mediated recombination through HMGB1/2 complex-induced 23-RSS DNA bending, showed no significant effect on recombination efficiency in our system (Figure S2). Using PiggyBac transposon, we established stable HEK293T clones with 1.27**–**2.31 antibody gene copies/cell (qPCR-verified, Figures S3A-C). Systematic optimization, combining RAG2 truncation mutants, mRNA delivery, and stable integration, boosted recombination efficiency 34.1-fold (1.32% to 45.01%, Figures S3D-F).

### 2.2. RSS-guided reprogramming enables comprehensive CDR1/2 diversification

To overcome the natural constraint limiting length variation to CDR3 in V(D)J recombination, we independently engineered modular 12/23-RSS elements into three CDR regions (CDR1/CDR2/CDR3) of HBsAg antibodies (**Figure 3A**). This synthetic redesign establishes V(D)J-like remodeling capability across the entire antigen-binding site, extending recombination-mediated diversification beyond CDR3 to regions (CDR1/CDR2) that normally undergo only SHM-mediated variation.

**Figure 3.**
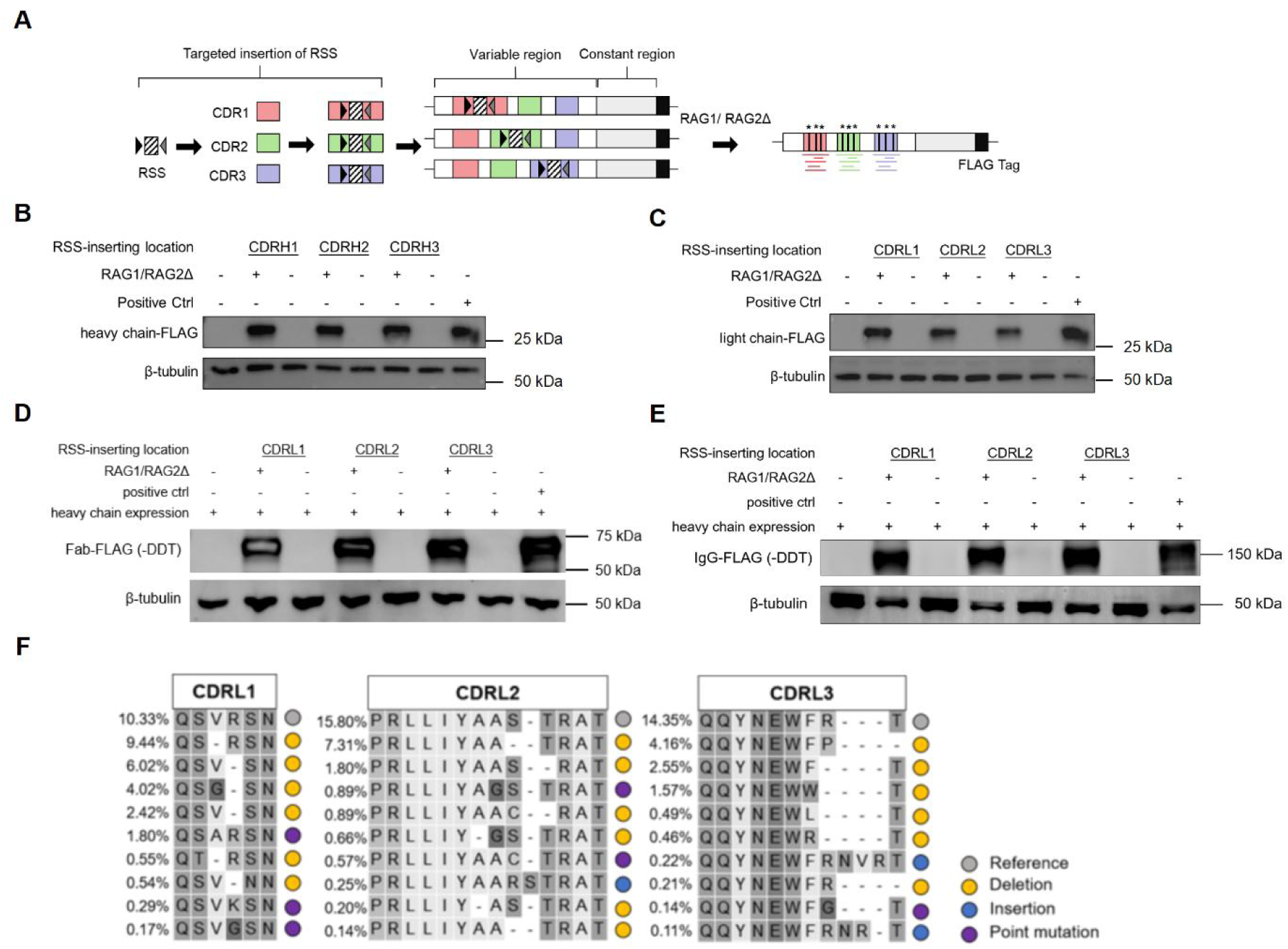
RSS-guided reprogramming of antibody genes. (A) Schematic overview of the antibody gene–reprogramming strategy, in which engineered recombination signal sequences (RSSs) direct RAG-mediated insertions into CDR1, CDR2, and CDR3. (B–C) Western blot analysis of heavy-chain (B) and light-chain (C) expression of Fab following targeted reprogramming of each CDR. Positive control: the correctly expressed Fab heavy chain(B)/light chain(C)-encoding sequence. (D) Assembly and expression of light chain-reprogrammed Fab. Analyzed by non-reducing SDS–PAGE and Western blot. Samples included the reprogrammed light chain of Fab co-expressed with its heavy chain. The parental Fab served as the positive control. (E) Assembly and expression of light chain-reprogrammed IgG. Analyzed by non-reducing SDS–PAGE and Western blot. Samples included the reprogrammed light chain of IgG co-expressed with its heavy chain. The parental IgG served as the positive control. (F) Top 10 most abundant amino acid sequences (by NGS read %) for CDRL1, CDRL2, and CDRL3 from HBsAg-binding antibodies after ARESEC rearrangement (B), annotated for mutation type (gray: reference; yellow: deletion; blue: insertion; purple: substitution).

Functional validation in HEK293T cells demonstrated that RAG1/2 mediated recombination at all engineered CDR regions while maintaining intact immunoglobulin structural integrity. Western blot analysis confirmed efficient expression of properly reprogrammed heavy chain (**Figures 3B**) and light chain (**Figures 3C**). Non-reducing SDS-PAGE with immunoblotting verified correct assembly of the reprogrammed chains into disulfide-linked Fab fragments (**Figure 3D**) and intact full-length IgG molecules (**Figure 3E**). Our mammalian display platforms achieved efficient surface localization of Fab and IgG (Figure S4A-C) with 27.8**–**28.2% of cells showing functional antigen binding. Combining ARESEC-mediated light chain reprogramming with subsequent light-heavy chain assembly, approximately 5% of cells still exhibited antigen-binding capability. This system, validated by immunofluorescence co-localization and flow cytometry (Figure S4A-C), enables high-throughput screening of structurally intact reprogrammed antibodies. These results establish that our RSS-mediated CDR reprogramming platform in non-lymphoid cell preserves native antibody folding and assembly pathways, including proper inter-chain disulfide bond formation and correct quaternary structure organization.

High-throughput sequencing of reprogrammed CDRL1, CDRL2, and CDRL3 revealed concentrated mutation hotspots within ±10 bp of recombination junctions (Figure S5). Deletion mutations predominated, particularly within 5 bp of cleavage sites where they surpassed 10% frequency. Analysis of in-frame sequences revealed that, at the protein level, only 10.33% (CDRL1), 15.8% (CDRL2), and 14.35% (CDRL3) of rearranged reads precisely reconstituted the parental antibody sequences, while a wide spectrum of mutations (predominantly amino acid deletions, followed by insertions and point mutation) was abundantly acquired (**Figure 3F**). These results demonstrate that RAG1/2-mediated recombination efficiently generates substantial junctional diversity near cleavage sites while maintaining structural integrity.

### 2.3. TdT Enhances Diversity in ARESEC-Reprogrammed Antibodies

Terminal deoxynucleotidyl transferase (TdT) was co-expressed with RAG1/2 in HEK293T cells to introduce non-templated nucleotides insertions (N-regions)^[4]^ at V(D)J junctions (**Figure 4A**). While maintaining recombination efficiency (**Figure 4B**), TdT increased unique sequences in CDRL1, CDRL2, and CDRL3 following ARESEC reprogramming by 41.5%-86.3% (**Figure 4C**). CDR length distributions shifted, with TdT-expressing variants (red) exhibiting greater sequence diversity across lengths than controls (blue) (**Figure 4D**). Although TdT expression did not alter the predominance of deletion mutations, it significantly increased the frequency of all mutation types. Quantitative analysis revealed that TdT expression specifically elevated insertion mutation frequencies within the ±5 bp region adjacent to RAG-mediated cleavage sites (**Figure 4E**). These results demonstrate that exogenous TdT enhances CDR sequence and length variability while maintaining overall recombination efficiency.

**Figure 4.**
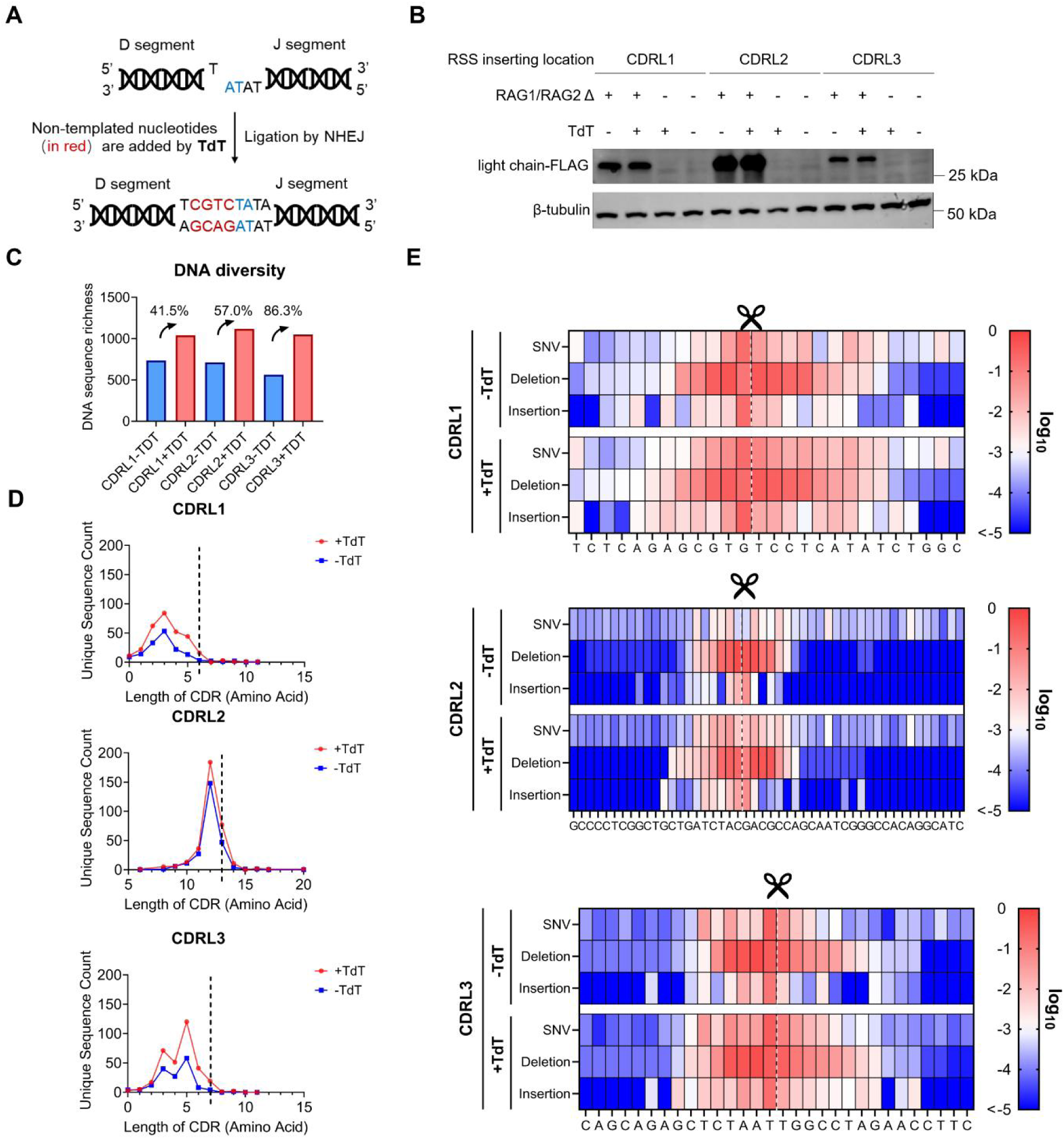
Optimization of antibody repertoire sequence diversity mediated by TdT. (A) Schematic diagram of the mechanism of TdT in V(D)J recombination. (B) Western blot analysis verifying the effect of TdT expression on the production of RAG1/2-mediated rearrangement products. (C) Impact of TdT expression on overall library diversity after ARESEC-mediated recombination. The richness (number of unique sequences) in DNA-based rearranged antibody libraries is compared between TdT-expressing (red) and control (blue) conditions. (D) Distribution of unique sequence counts by CDR length. Red, +TdT; Blue, -TdT; Dashed line, CDR length of the parental antibody. (E) Impact of TdT expression on junctional diversity following antibody gene rearrangement. This sequencing analysis depicts the distribution and frequency of nucleotide mutation types at each position in CDRL1, CDRL2, and CDRL3 following rearrangement by ARESEC, with or without TdT. The heatmap shows log₁₀-transformed values of mutation frequency. Scissors and dashed lines indicate RAG1/2 recognition and cleavage sites.

### 2.4. ARESEC-mediated Engineering of the clinical antibody Nivolumab

To evaluate ARESEC’s capability in antibody affinity maturation and structural reprogramming, we applied it to engineer Nivolumab, a clinical anti-PD-1 antibody^[51]^. Structural analysis revealed that Nivolumab (a PD-1 inhibitor with analogous binding topology) engages the PD-1 N-terminal loop region (BC/CD/FG loops), sterically hindering PD-L1 interaction through a “molecular wedge” mechanism (**Figure 5A-B**)^[52]^. Notably, Nivolumab’s light chain establishes merely 4 PD-1 contacts versus Pembrolizumab’s 13^[53]^, suggesting suboptimal light chain engagement and presenting a prime target for ARESEC-mediated light chain optimization.

**Figure 5.**
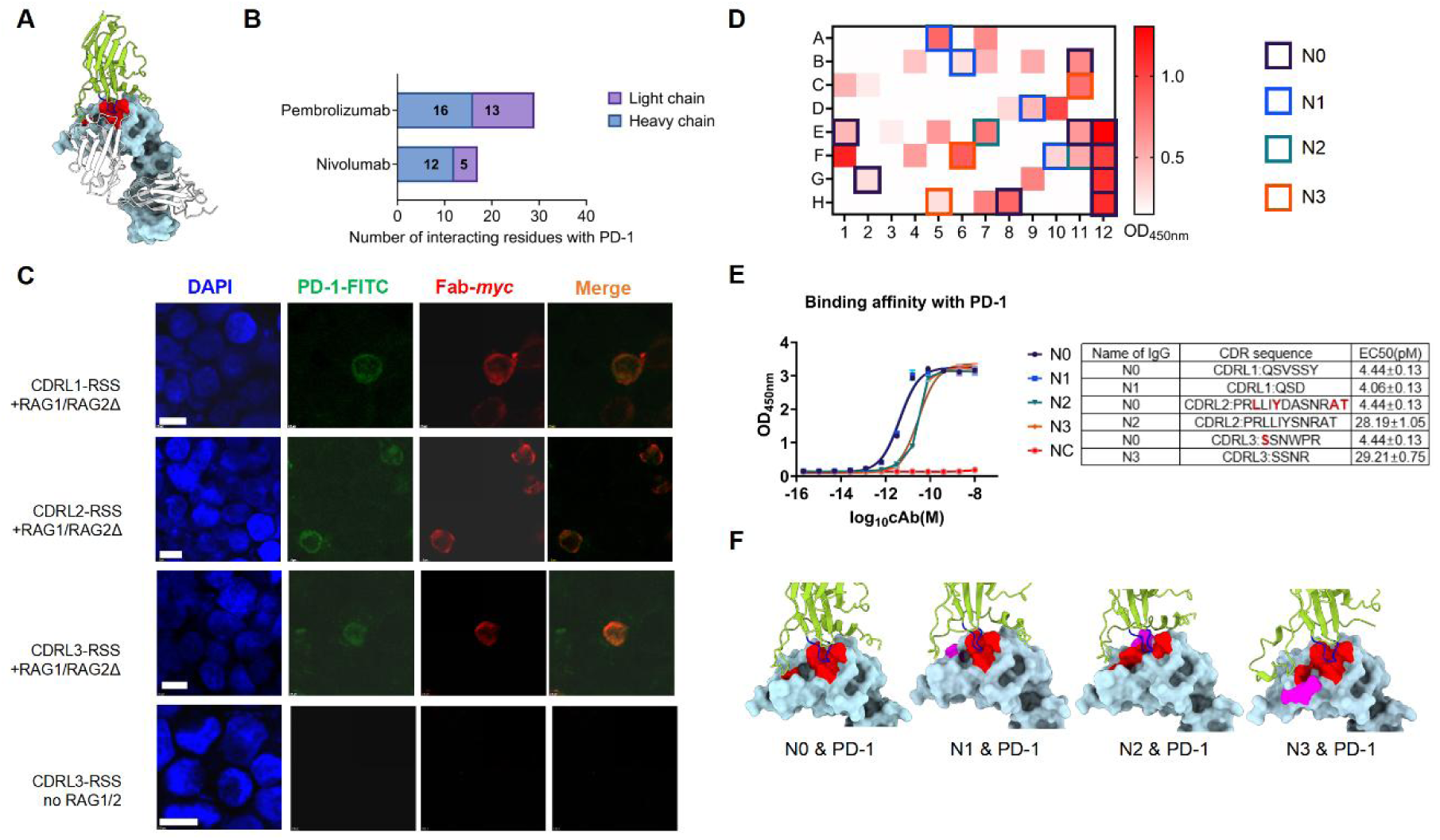
Reprogramming and screening of the PD-1 antibody Nivolumab using the ARESEC system. (A) Schematic representation of the PD-1–Nivolumab complex. PD-1 is shown in surface representation (yellow-green), and the heavy and light chains of the Nivolumab Fab are shown in cartoon representation (white and light blue, respectively). The binding interface is highlighted, with the PD-1 contact loop shown in blue and Fab paratope residues shown in red. (B) Statistical analysis of the interaction sites between PD-1 and Nivolumab or Pembrolizumab respectively. Data for the interaction sites were obtained from previous reports. (C) Immunofluorescence staining showing antibody display on the cell surface. Red fluorescence indicates the myc tag labeled with AF700-conjugated antibody, green fluorescence shows FITC-conjugated PD-1 antigen, and blue represents DAPI-stained nuclei. Scale bar:5 μm. (D) Primary cell ELISA screening of flow-sorted clones and their PD-1 binding capacity relative to parental Nivolumab. The intensity of the red color is proportional to the OD450nm reading, representing a combined measure of antibody affinity and expression level. A total of 28 candidate high-affinity clones were obtained, belonging to 10 distinct sequences. Among these, the three variants (N1, N2, N3) with the highest average absorbance at 450 nm, along with the wild-type (N0), were selected for expression (indicated by boxes). (E) ELISA-based measurement of the binding affinity of purified Nivolumab and its variants N1, N2, N3 for PD-1. Residues directly interacting with PD-1 are in red. (F) Structural models of Nivolumab (N0) and mutant Fabs (N1-N3) in complex with PD-1. Close-up of the binding interface: PD-1 contact loop (blue), Fab paratope residues (red), and mutation sites (magenta). Models predicted by AlphaFold3; interface annotations based on published complexes (PDB: 5WT9/5GGR).

Analysis of reprogrammed Nivolumab light chains showed 23**–**33% recombination efficiency (PCR) and proper protein expression (Western blot); these recombination efficiencies proved consistent with those from our previous validation systems, confirming the robustness and reproducibility of the ARESEC platform. High-throughput sequencing identified deletion-rich variants (<13% parental sequences) with CDR-specific mutation biases (Figure S6A-C). Functional validation by immuno-fluorescence (**Figure 5C**) confirmed membrane-displayed antibodies (red, HC-myc-AF700) co-localized with PD-1 antigen (green, PD-1-FITC), indicating proper antibody folding and antigen-binding capability in HEK293T cells. Two rounds of FACS and ELISA screening yielded 28 high-affinity monoclonal candidates, among which 10 distinct antibody sequences were identified by sequencing (**Figure 5D**). We selected three top variants (N1-N3) from initial screening, along with parental Nivolumab (N0), for IgG4 expression and purification in HEK293F cells (Figure S7). ELISA-based affinity assays revealed that N1 (CDRL1: QSVSSY→QSD), despite comprising <0.2% of the library, exhibited a modest 1.1-fold affinity gain (EC50 = 4.06±0.13 pM) over N0 (4.44±0.13 pM) (**Figure 5E**). Although spatially distal to the paratope-epitope interface (**Figure 5F**), the QSD mutation subtly enhanced binding affinity. In contrast, N2 and N3 exhibited significant affinity reductions (6.3-fold and 6.8-fold, respectively) due to CDR loop deletions proximal to the binding interface: N2 lacks CDRL2 residues “DA” near the interfacial region, while N3’s truncated CDRL3 (“SSNWPR” → “SSNR”) abuts the critical antigenic residue S91. These results demonstrate that non-paratopic mutations-even those not directly contacting the antigen-can allosterically modulate binding affinity, underscoring ARESEC’s unique capability to uncover such non-canonical regulatory variants.

### 2.5. ARESEC platform Enhances Durvalumab’s Binding Affinity

To evaluate the system’s versatility in therapeutic antibody engineering, we applied ARESEC to reprogram Durvalumab, a high affinity anti-PD-L1 IgG1 (**Figure 6A**) that blocks PD-L1/PD-1 interactions to restore T-cell immunity^[54] [55]^. RAG1/2-mediated recombination in Durvalumab light chain CDRs was successfully achieved, with efficiencies of 38.24% (CDRL1), 47.27% (CDRL2), and 35.23% (CDRL3) as confirmed by PCR and Western blot analysis (Figure S8A-B). High-throughput sequencing revealed over 1000 unique sequences per CDR, with deletion mutations (>10% frequency) predominantly localized within ±10 bp of RAG1/2 cleavage sites (Figure S8C). Following cell pool expansion and screening (**Figure 6B**), we selected two clones (Dur1 and Dur2) with enhanced binding for sequencing, IgG expression, and purification (**Figure 6C**). ELISA affinity measurements demonstrated that Dur1 (CDRL2: PRLLIYDASSRAT→PRLLYDASSRAT, 2.31 ± 0.02 pM) achieved a 2.01-fold affinity improvement over wild-type Durvalumab (Dur0, 4.65 ± 0.16 pM), while Dur2 (CDRL2: PRLLIYDASSRAT → PRLYDASSRAT, 3.03 ± 0.10 pM) showed a 1.53-fold improvement (**Figure 6D**). Structural modeling indicated that both variants contained CDRL2 deletions distal to CDRL3 ’ s direct interaction sites (**Figure 6E**). These results demonstrate ARESEC’s capacity for efficient CDR remodeling while preserving structural integrity, and establish its clinical potential through successful optimization of a therapeutic antibody.

**Figure 6.**
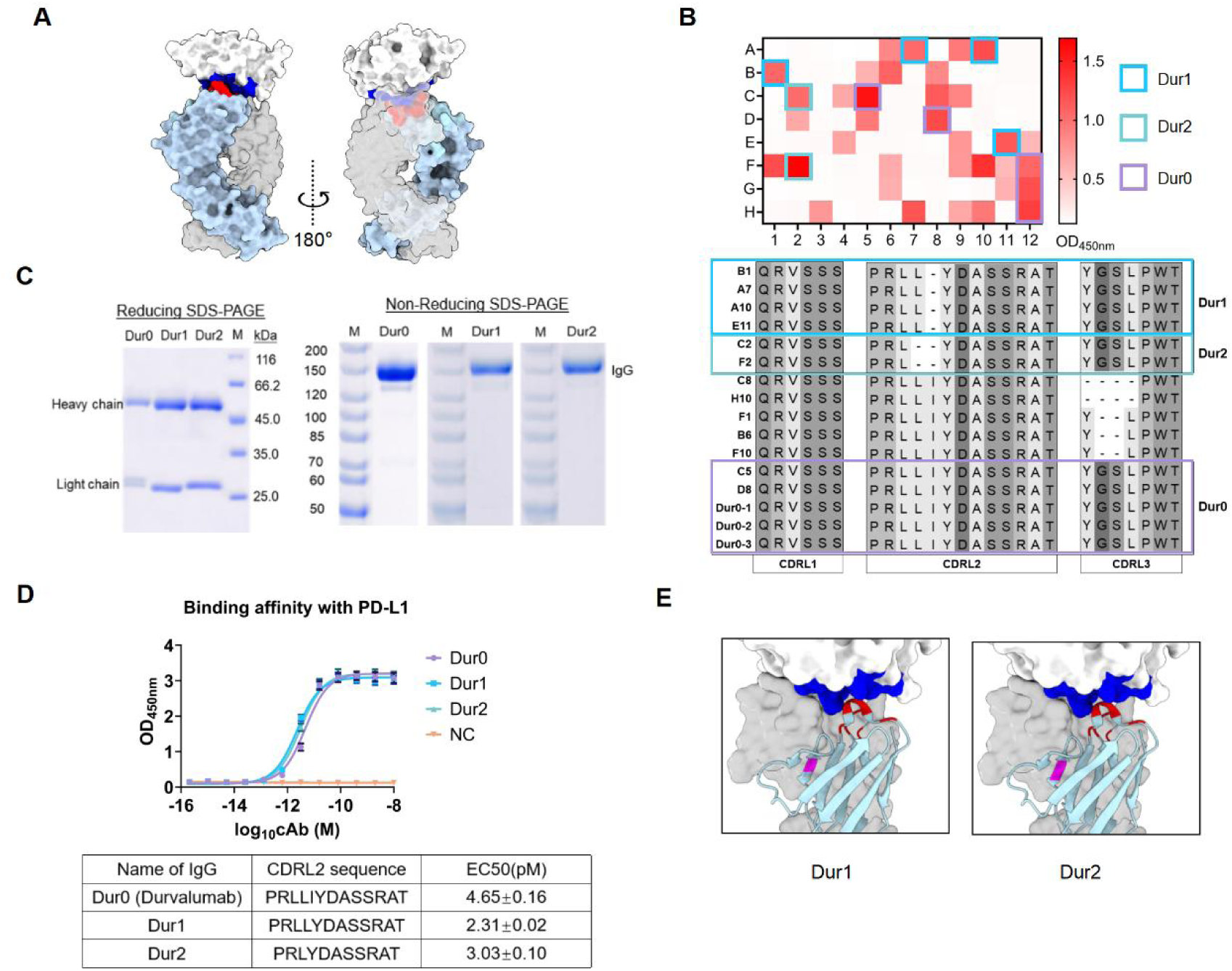
Reprogramming and screening of the PD-L1 antibody Durvalumab using the ARESEC system. (A) Structural model of the PD-L1/Durvalumab complex (based on PDB 5X8M). PD-L1, the heavy chain of Durvalumab, and the light chain of Durvalumab are colored white, gray, and light blue, respectively. Residues participating in direct interactions between the light chain of the Durvalumab Fab and PD-L1 are indicated in red and blue. (B) Top, primary cell ELISA screening of flow-sorted clones and their PD-L1 binding capacity relative to parental Durvalumab. Bottom, CDR sequences of the selected monoclonal antibodies from the screening. Four distinct antibodies were identified, among which the 2 variants exhibiting the highest absorbance were selected for subsequent expression and purification. (C) SDS-PAGE under reducing/non-reducing conditions. Lane M: Protein molecular weight marker; Lanes Dur0-Dur2: Purified antibody samples Dur0-Dur2, showing bands at approximately 50 kDa (Heavy Chain) ,25 kDa (Light Chain) and 150 kDa (IgG). (D) ELISA measurement of the binding affinity of Durvalumab (Dur0) and its variants Dur1 and Dur2 to PD-L1. (E) Structural model of mutation sites in Dur1/Dur2 based on the complex structure (PDB: 5X8M). PD-L1, the heavy chain of Durvalumab, and the light chain of Durvalumab are colored white, gray, and light blue, respectively. Residues involved in direct interactions between the light chain of the Durvalumab Fab and PD-L1 are indicated in red and blue. Deletion sites specific to Dur1 and Dur2 are highlighted in magenta.

## 3. Discussion

V(D)J recombination has traditionally been considered lymphocyte-specific, but recent advances enable its reconstitution in microbial cells, offering new possibilities for antibody engineering^[56]^. The pioneering work of Cazier *et al.* achieved V(D)J-like recombination in *S. cerevisiae*, establishing conceptual groundwork for extra-lymphoid recombination^[57]^. Yet this yeast platform suffers from low recombination efficiency (∼1%) and lacks natural junctional diversity due to homology-driven repair. Unlike yeast-based systems, the ARESEC developed in this study leverages endogenous human TdT and NHEJ repair machinery to generate natural junctional diversity (N-additions and hairpin resolution). The platform’s physiological recombination efficiency (>45%) and mammalian cell compatibility allow for direct selection of functional variants against native antigens. Notably, the ARESEC platform extends V(D)J recombination to all three CDRs, overcoming 500 million years of evolutionary constraint that limited natural diversification primarily to CDR3. This system maintains robust structural integrity, validated through surface display and functional assays.

The integrated technological approach delivers several transformative advantages for therapeutic antibody development. First, ARESEC overcomes the limitations of conventional approaches by enabling comprehensive antibody reprogramming beyond CDR3-focused modifications. Second, these capabilities were validated through the successful affinity maturation of the clinical anti-PD-L1 antibody Durvalumab, achieving a 2.01-fold improvement in binding affinity (KD from 4.65 pM to 2.31 pM) through engineered CDR2 modifications while preserving structural stability. Collectively, these results demonstrate ARESEC’s capacity to outperform conventional antibody optimization methods by enabling efficient, multi-CDR remodeling with high functional retention.

The RAG recombinase system exhibits unique advantages over CRISPR-Cas9 for *in vivo* antibody diversification^[57]^. Its operational superiority stems from three mechanistic features: (1) Precision targeting: RSS-guided cleavage (12/23 rule) ensures coordinated double-strand breaks at paired gene segments; (2) Structural fidelity: Native hairpin formation enables physiological processing of coding joints; and (3) Combinatorial efficiency: Single-step generation of diverse V-J combinations (vs. CRISPR’s requirement for synchronous multi-guide cleavage). Through ARESEC implementation, we engineered V-J rearrangement by coupling 12-RSS to ends of two V-gene and 23-RSS to J-segments, successfully producing combinatorial antibody diversity (unpublished data). This multiplexed reprogramming would demand prohibitively complex coordination of ≥ 3 concurrent Cas9 cleavages to achieve comparable repertoire breadth. Thus, the RAG recombinase system is inherently more suitable than CRISPR-Cas9 for orchestrating recombination across multicopy V, D, and J gene segments.

The current ARESEC system exhibits biased diversification favoring nucleotide deletions over balanced P/N-nucleotide additions, despite confirmed TdT activity^[58]^. This imbalance likely stems from suboptimal coordination between RAG cleavage and DNA repair pathways in HEK293T cells, resulting in incomplete N-region expansion and modest library diversity. Attempts to enhance diversity through Artemis over-expression proved unsuccessful, suggesting broader NHEJ machinery limitations in this non-lymphoid context^[59]^. Future development should focus on co-expressing core NHEJ components (DNA-PKcs, Ku70/80, XLF) to improve repair coordination, optimizing TdT expression for enhanced N-nucleotide insertion, and scaling library generation capacity. These refinements will be crucial for achieving physiological diversification patterns and maximizing functional repertoire generation.

In summary, ARESEC represents a novel synthetic biology platform that fundamentally redefines antibody diversification by liberating V(D)J recombination from its evolutionary constraints. This system provides a valuable toolkit that not only advances fundamental antibody engineering but also holds strong potential for accelerating therapeutic discovery.

## 4. Experimental Section

### 4.1. Cells and Strains

HEK293T and HEK293F cell lines were obtained from the laboratory of Professor Yuhang Zhang and routinely authenticated by short tandem repeat (STR) profiling. All cell cultures were confirmed to be free of mycoplasma contamination using a PCR-based detection kit (Beyotime, C0301S). HEK293T cells were maintained in Dulbecco’s modified Eagle’s medium (DMEM, high glucose, Gibco, 11995065) supplemented with 10% (v/v) fetal bovine serum (FBS, Gibco, 10099141) and 1% (v/v) penicillin-streptomycin (Gibco, 15140122) at 37°C in a 5% CO₂ humidified atmosphere. Cells were passaged every 2-3 days using 0.25% Trypsin-EDTA (Gibco, 25200056) and routinely monitored for morphology and confluence. This adherent cell line served as the primary platform for genomic integration, RAG-mediated reprogramming, mammalian surface display, and functional screening. HEK293F suspension cells were cultured in FreeStyle™ 293 Expression Medium (Thermo Fisher Scientific, 12338018) in polycarbonate Erlenmeyer flasks with vented caps, incubated at 37°C under 8% CO₂ with constant orbital shaking at 120 rpm. Cells were passaged every 3-4 days to maintain a density between 0.5×10^6^ and 3.0×10^6^ cells/mL and were used exclusively for transient expression and large-scale production of full-length IgG antibodies.

*Escherichia coli* DH5α chemically competent cells (Weidi Bio, DL1001) were used for plasmid construction and propagation. Cells were cultured in LB medium supplemented with appropriate antibiotic at 37°C with shaking at 220 rpm. Single colonies were isolated on LB agar plates containing the appropriate antibiotic after transformation and used for plasmid DNA preparation.

### 4.2. Generation of antibody expression constructs designed for programmed recombination

Based on the V(D)J recombination mechanism, a modular recombination signal system containing 12-RSS and 23-RSS sequences was designed. A complete recombination control unit was formed by inserting an SV40 poly(A) signal sequence between the two RSS sequences. These modular elements were individually inserted into the CDR1, CDR2, and CDR3 regions of a mammalian codon-optimized antibody light chain gene (synthesized by GenScript) via molecular cloning. The final gene expression cassette included a PGK promoter-driven puromycin resistance gene, a C-terminal 3×FLAG tag sequence on the light chain, and a P2A self-cleaving peptide-linked blasticidin S deaminase gene. All functional modules were assembled into the pcDNA3.1 vector backbone using a seamless cloning kit (Vazyme, C115-02) with a 1-hour reaction at 50°C, resulting in the reprogrammable antibody gene cassette. The recombinant plasmid was transformed into *E. coli* DH5α competent cells via heat shock, selected on LB plates containing appropriate antibiotics, and validated by restriction enzyme digestion after monoclonal expansion.

### 4.3. PiggyBac Vector Construction and Stable Cell Line Establishment

The PiggyBac transposon system was employed for genomic integration of the antibody gene ^[60]^. Vector backbone and light chain insert fragments (with ∼20 bp overlaps) were amplified by PCR and assembled using the seamless cloning kit (Vazyme, C115-02) incubated at 50°C for 1 hour. The recombinant plasmid was transformed into *E. coli* DH5α competent cells via heat shock, selected on LB plates containing ampicillin (100 μg/mL), and validated by restriction enzyme digestion after monoclonal expansion. HEK293T cells were seeded in 6-well plates at 1×10^6^ cells/well. Transfection was performed at 70% confluency using a mixture of 1 μg PiggyBac transposon vector and 1.5 μg transposase expression vector, complexed with 4 μL Lipo8000 transfection reagent (Beyotime) in 250 μL serum-free DMEM. After 15 minutes at room temperature, the complexes were added to the cells. The medium was replaced with complete medium 6 hours post-transfection, and puromycin selection (1.5 μg/mL) began 48 hours later, continuing for 7-14 days to establish stable cell lines.

### 4.4. RAG1/2-Mediated Antibody Gene Reprogramming

Cells stably integrated with the antibody light chain gene were seeded in 10 cm dishes at 5-8×10^6^ cells/dish. At 80% confluency, transfection was performed with 15 μg RAG1 expression plasmid, 15 μg RAG2(1-361) truncated mutant mRNA, and 30 μL Lipofectamine 3000 reagent (Thermo Fisher Scientific, L3000015) in 1 mL Opti-MEM (Gibco, 31985070). After 6 hours, the medium was replaced with fresh complete medium. Blasticidin S (10 μg/mL) selection started 48 hours post-transfection and continued for 7-14 days to obtain the reprogrammed cell population.

### 4.5. Gene Copy Number Quantification

Genomic DNA was extracted using a DNA extraction kit (Beyotime, D0063). Plasmid DNA containing the target gene fragment served as the quantitative standard, serially diluted (10^3^-10^9^ copies/μL). The qPCR reaction contained 100 ng template DNA, 0.3 μM specific primers, and 10 μL SYBR Green Master Mix in a 20 μL total volume. The program was: 95°C for 5 min; 40 cycles of 95°C for 10 sec, 60°C for 30 sec, 72°C for 30 sec. Absolute copy numbers were calculated from the standard curve^[61]^.

### 4.6. PCR Analysis of Reprogramming Efficiency

Specific primers flanking the modular RSS insertion sites were used for amplification with KOD One^TM^ PCR Master Mix. The 50 μL reaction contained 1 μg template DNA and 0.3 μM each primer. The PCR program was: 94°C for 2 min; 25 cycles of 98°C for 10 sec, 58°C for 10 sec, 68°C for 10 sec; final extension at 68°C for 3 min. Products were analyzed by 1% agarose gel electrophoresis^[62]^.

### 4.7. Western Blot Analysis of Protein Expression

Total protein was extracted using RIPA lysis buffer with protease inhibitors. After 10 minutes on ice and centrifugation at 12,000 rpm for 5 minutes, the supernatant was collected. Protein concentration was determined by BCA assay and normalized. 50 μg of protein per sample was separated by SDS-PAGE, transferred to PVDF membrane at 280 mA for 75 minutes, blocked with 5% BSA, and sequentially incubated with anti-FLAG primary antibody (1:5000) and HRP-conjugated secondary antibody (1:6000). Signals were detected using ECL substrate. The bands detected by E-blot typically migrate at a slightly higher molecular weight than theoretically predicted. This shift is due to residual amino acids that remain attached following incomplete cleavage of the P2A peptide.

### 4.8. NGS Analysis of Sequence Diversity

Genomic DNA was extracted from approximately 5×10⁶ reprogrammed HEK293T cells using the DNA extraction kit (Beyotime, D0063). A two-step PCR strategy^[45]^ was employed for library preparation: the first PCR amplified the variable regions flanking the engineered RSS sites with gene-specific primers using Q5 High-Fidelity DNA Polymerase (16 cycles), followed by a second PCR to append full-length Illumina adapters and sample indices. The purified libraries were quantified, pooled, and sequenced on the Illumina MiSeq platform using 2×250 bp paired-end chemistry. Raw sequencing reads were subjected to quality filtering with fastp to remove adapters and low-quality bases (Q30 ≥ 94.66%). Paired-end reads were merged using FLASH (efficiency > 98.6%), and the resulting contigs were aligned to the reference sequence via Clustal W. Polymorphism profiling and variant calling were performed based on multiple sequence alignment generated by MAFFT, with Single Nucleotide Variant (SNV) and insertion–deletion variants (Indels) identified and annotated by mutation type, allele frequency, and read support. To minimize sequencing-derived diversity bias, all processed sequences were systematically filtered against reference control samples (parental sequences) to exclude background noise and systematic artifacts prior to downstream analysis. Diversity metrics were calculated from DNA sequences supported by more than 100 reads, and positional amino acid frequencies were analyzed using R.

### 4.9. Co-expression of Terminal Deoxynucleotidyl Transferase (TdT) for Antibody Sequence Diversification

To enhance junctional diversity during antibody gene assembly, terminal deoxynucleotidyl transferase (TdT) was co-expressed with the core RAG1/2 recombinase components. Using the Nivolumab sequence as the parental antibody backbone, a codon-optimized human TdT gene (synthesized by GenScript) under the control of a CMV promoter was cloned into an expression vector. This TdT construct was co-transfected with the ARESEC-reprogramming plasmids into HEK293T cells. Control transfections were performed in parallel using the same RAG1/2 and ARESEC plasmids without the TdT expression vector.

### 4.10. Mammalian Cell Surface Display

For antibody surface display, the CH1 domain C-terminus of the Fab fragment was fused to the PDGFR transmembrane domain. Fixed cells (4% PFA) were sequentially incubated with Myc tag (Alexa Fluor 700) antibody (1:1000) and FITC-labeled antigen (1:100) in the dark, followed by DAPI (1 μg/mL) nuclear staining. Localization was observed by confocal microscopy^[63] [64]^.

### 4.11. Flow Cytometry and Cell Sorting

Flow cytometry analysis and fluorescence-activated cell sorting (FACS) were performed using Cytoflex (Beckman) and BD FACS Aria^TM^ III instruments (BD Biosciences), respectively. To quantify the RAG recombination efficiency, cells were harvested 48 h post-transfection, detached with trypsin, and washed three times with cold phosphate-buffered saline (PBS). The proportion of EGFP-positive cells was determined using the Cytoflex flow cytometer. For antibody surface-display screening, cells expressing Fab fragments were similarly collected and washed with cold PBS. Cell pellets containing 5 × 10⁵ cells were resuspended in 50 µL of staining buffer containing FITC- or Cy3-conjugated antigen (1:100 dilution) and AF700-labeled anti-myc antibody (1:100 dilution). After incubation on ice for 30 min in the dark, cells were washed twice with PBS and resuspended in PBS for analysis or sorting. All centrifugation steps were carried out at 250 × g for 5 min at 4 °C. For experiments with larger cell numbers, staining volumes were scaled proportionally to maintain a ratio of 100 µL labeling mix per 1 × 10^6^ cells. Unstained cells and cells expressing only the heavy chain or light chain served as negative controls. Double-positive populations were identified by sequential gating and isolated by FACS for downstream validation^[65]^.

### 4.12. ELISA Using Cell Lysates for Membrane Antibody Detection

Cells were gently washed with cold PBS and lysed on ice with RIPA buffer containing protease inhibitors. After centrifugation at 12,000 g for 15 minutes at 4°C, the supernatant (total cell protein lysate) was collected. The antigen (PD-1, His-tagged recombinant human protein, GenScript, Z03424; PD-L1, His-tagged recombinant human protein, GenScript, Z03425) was diluted to a working concentration of 1 µg/mL in coating buffer (0.05 M carbonate-bicarbonate buffer, pH 9.6, Sigma Aldrich, C3041). A volume of 100 µL per well was added to a 96-well high-binding polystyrene microplate (Corning, 9018) and incubated overnight at 4 °C. Following three washes with washing buffer (PBS containing 0.05% Tween-20, PBST), the plate was blocked with 200 µL/well of blocking buffer (5% w/v non-fat dry milk in PBST) for 1 h at 37 °C. Purified antibodies were serially diluted in antibody dilution buffer (1% BSA in PBST) and added to the plate (100 µL/well) in triplicate, followed by incubation at 37 °C for 1.5 h. After washing, the plate was incubated with horseradish peroxidase (HRP)-conjugated goat anti-human IgG secondary antibody for 1h at 37°C. Colorimetric development was performed by adding 100 µL/well of TMB substrate solution (Sigma Aldrich, T0440) and incubating for 15 min in the dark. The reaction was stopped with 50 µL/well of 2M sulfuric acid, and absorbance was measured at 450 nm using a microplate reader (BioTek Synergy 2) ^[67]^.

### 4.13. Antibody Expression and Purification

HEK293F cells in logarithmic growth phase (density 2×10^6^/mL) were co-transfected with heavy and light chain expression plasmids at a 1:1 ratio (total 30 μg DNA) using polyethylenimine (PEI). After 4 days of culture at 37°C with shaking, the supernatant was collected, clarified by centrifugation, and purified using Protein A/G affinity chromatography. The bound antibody was eluted with 0.1 M glycine (pH 3.0) and immediately neutralized with Tris-HCl (pH 8.0). Antibody concentration was determined by NanoDrop spectrophotometer: 1-2 μL sample was measured at 280 nm using an extinction coefficient of 1.4 (mg/mL) ^-1^cm^-1^; purity was assessed by size exclusion chromatography-HPLC using a biocompatible SEC column with phosphate buffer isocratic elution (flow rate 0.5-1.0 mL/min); 20-50 μg sample was injected and elution monitored at 280 nm; monomer purity and aggregate content were calculated from peak area integration ^[68]^.

### 4.14. Reducing and Non-Reducing SDS-PAGE

To evaluate antibody integrity and disulfide bond status, samples were prepared under reducing (2% SDS, 100 mM DTT, 95°C for 10 min) and non-reducing (SDS loading buffer without reductant, same heating) conditions. 20 μL samples (50 μg protein) alongside pre-stained markers were loaded onto 4-20% polyacrylamide gels and electrophoresed at 80-120 V for ∼90 minutes in Tris-glycine buffer containing 0.1% SDS. Gels were stained with Coomassie Brilliant Blue R-250, destained, imaged, and band patterns analyzed to assess assembly integrity and detect aggregation or degradation ^[69]^.

### 4.15. Antigen-Antibody Affinity ELISA

The antigen (PD-1, His-tagged recombinant human protein, GenScript, Z03424; PD-L1, His-tagged recombinant human protein, GenScript, Z03425) was diluted to a working concentration of 1 µg/mL in coating buffer (0.05 M carbonate-bicarbonate buffer, pH 9.6, Sigma Aldrich, C3041). A volume of 100 µL per well was added to a 96-well high-binding polystyrene microplate (Corning, 9018) and incubated overnight at 4 °C. Following three washes with washing buffer (PBS containing 0.05% Tween-20, PBST), the plate was blocked with 200 µL/well of blocking buffer (5% w/v non-fat dry milk in PBST) for 1 h at 37 °C. Purified antibodies were serially diluted in antibody dilution buffer (1% BSA in PBST) and added to the plate (100 µL/well) in triplicate, followed by incubation at 37 °C for 1.5 h. After washing, the plate was incubated with horseradish peroxidase (HRP)-conjugated goat anti-human IgG secondary antibody for 1h at 37°C. Colorimetric development was performed by adding 100 µL/well of TMB substrate solution (Sigma Aldrich, T0440) and incubating for 15 min in the dark. The reaction was stopped with 50 µL/well of 2 M sulfuric acid, and absorbance was measured at 450 nm using a microplate reader (BioTek Synergy 2)^[70]^.

### 4.16. Statistical Analysis

Statistical analyses were performed using GraphPad Prism 10.1.2. Continuous data are presented as mean ± standard deviation (SD). Differences between two independent groups were analyzed using an unpaired two-tailed Student’s t-test for normally distributed data. Comparisons among three or more groups were conducted by one-way or two-way ANOVA. Statistical significance is indicated as follows: *p< 0.05, **p< 0.01, ***p< 0.001, and ****p< 0.0001; “n.s.” denotes non-significance (p≥ 0.05).

## Acknowledgements

We sincerely acknowledge the Instrument-Sharing Platform at the School of Life Sciences and Biotechnology, Shanghai Jiao Tong University for invaluable support. We also wish to express our gratitude to Haixia Jiang (SJTU) for her assistance with flow cytometry experiments, and to Feiyang Xue (SJTU) for his expert guidance in confocal microscopy. This research was supported by grants from National Key Research and Development Program of China (2023YFA0913700), Shanghai Municipal Science and Technology Major Project, and the Project of State Key Laboratory of Microbial Metabolism, Shanghai Jiao Tong University (SKLMMCX25_01).

## Data Availability Statement

Raw NGS sequencing data are available from the NCBI database under Sequence Read Archive (SRA) accession number PRJNA1373215, PRJNA1373465, PRJNA1373144 and PRJNA1375484.

---

**Figure S1.**
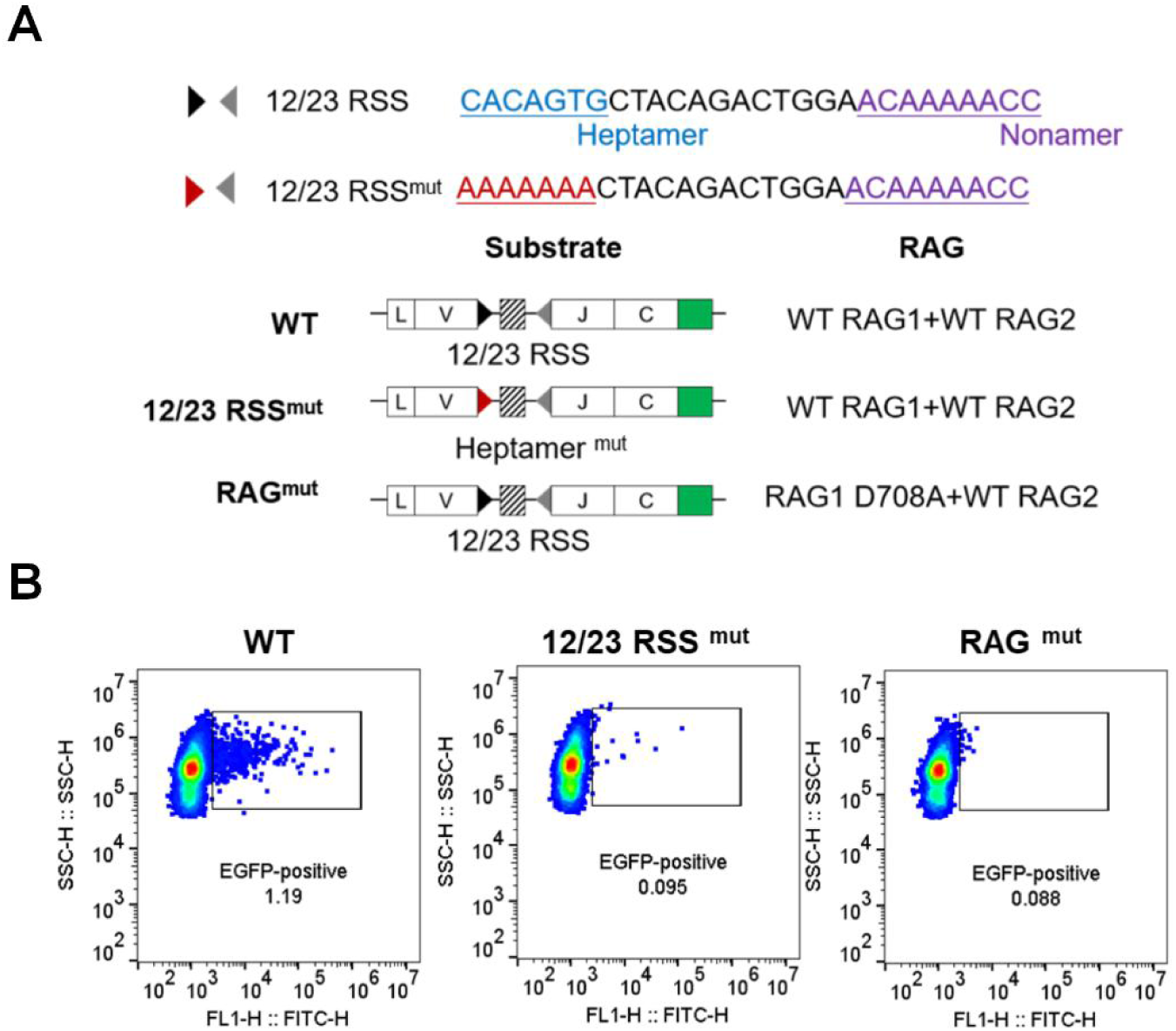
Defining the essential requirements for V-J recombination in HEK293T cells, related to Figure 2. (A) Schematic of mutant constructs assessing essential factors for V-J rearrangement in HEK293T cells. The Wild-Type (WT) group contains functional 12/23 RSS (heptamer in blue) and RAG1. The 12/23 RSS^mut^ group carries a heptamer mutated to AAAAAAA (red), disrupting RAG recognition. The RAG^mut^ group expresses a catalytically inactive RAG1 (D708A) to abrogate recombination activity. (B) Flow cytometric analysis of recombination efficiency. HEK293T cells were transfected with the EGFP reporter plasmid along with different RAG combinations: (i) wild-type RAG1/RAG2(WT), (ii) RSS heptamer mutant (12/23RSS^mut^), and (iii) RAG1 catalytic dead mutant (RAG^mut^).

**Figure S2.**
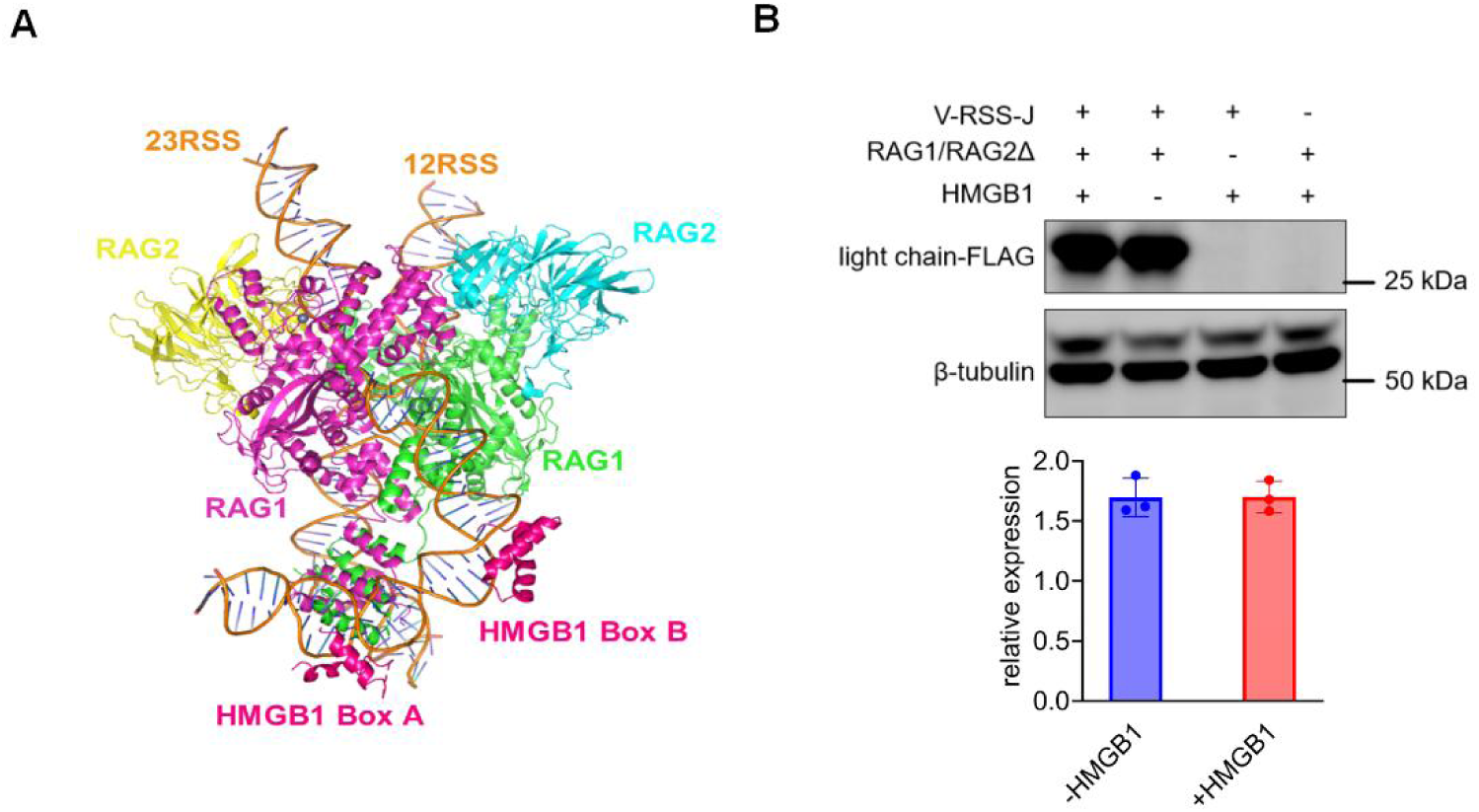
Assessment of HMGB1 co-expression on RAG-mediated recombination efficiency, related to Figure 2. (A) Atomic model of the RAG-RSS-HMGB1 complex (based on PDB:6CG0). (B) Western blot analysis of light-chain. HEK293T cells were co-transfected with RAG1/2 expression constructs alongside either an empty vector control or an HMGB1 expression plasmid. Recombination efficiency was evaluated by Western blot detecting FLAG. β-tubulin served as a control.

**Figure S3.**
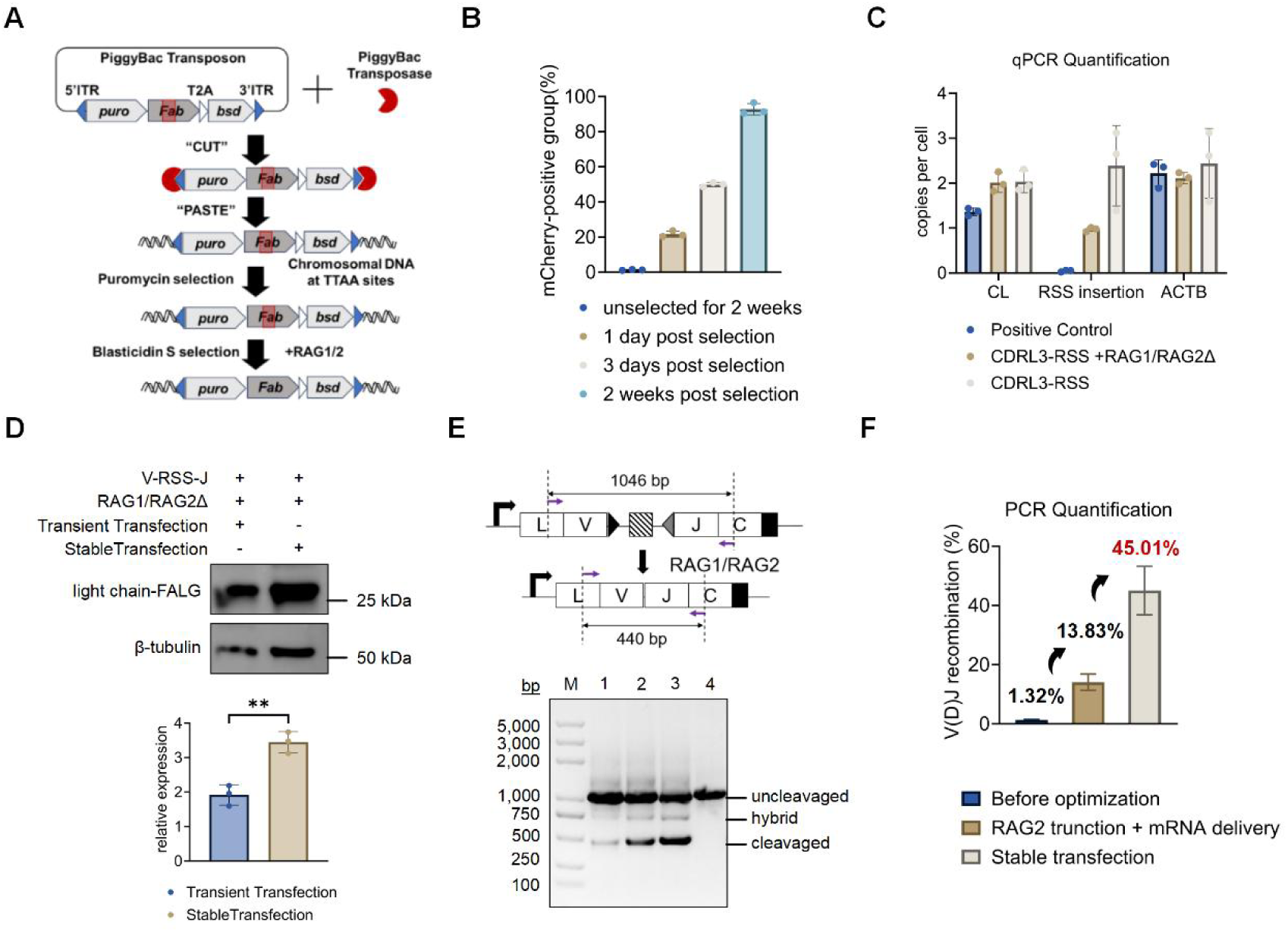
Construction and recombination of stable-integration HEK293T carrying antibody gene cassettes, related to Figure 2. (A) Schematic of the workflow for generating stable cell lines and subsequent antibody gene reprogramming. The diagram illustrates the construction and selection of stable cell lines using the piggyBac transposon system, followed by antibody gene reprogramming via co-transfection of a RAG1 expression plasmid and a truncated RAG2Δ mRNA. The final step involves enrichment of antibody-expressing cell populations by blasticidin S selection. (B) Analysis of antibody gene integration efficiency mediated by the piggyBac transposon system. Flow cytometry analysis showing the percentage of mCherry-positive HEK293T cells after co-transfection with a piggyBac-mCherry reporter plasmid and a transposase expression plasmid, followed by puromycin selection for varying durations. (C) qPCR analysis of antibody gene copy number in the HBSAb light chain-stable integrating HEK293T.Primers were designed specificly binding to constant region and RSS insertion of light chain coding gene. The absolute copy number per cell was calculated using a standard curve, with the ACTB gene serving as the internal reference control. (D) Western blot analysis of antibody gene rearrangement products in HEK293T cells. Target antibody genes were delivered through either transient plasmid transfection (lane 1) or stable cell line generation (lane 2), followed by co-expression of RAG1 and RAG2. (E) PCR-based assessment of RAG1/2-mediated antibody gene rearrangement under optimized conditions. The purple arrows indicate the locations of the primers. Lane M: DNA ladder; Lane 1: Pre-optimization control; Lane 2: Stage 1 optimization (RAG2 truncation mutant + mRNA delivery); Lane 3: Stage 2 optimization (stable transfection + antibiotic selection); Lane 4: Negative control (no RAG1/2). (F) Bar graph summarizing the optimized efficiency of RAG1/2-mediated antibody gene rearrangement, as quantified from the data shown in panel (**E**).

**Figure S4.**
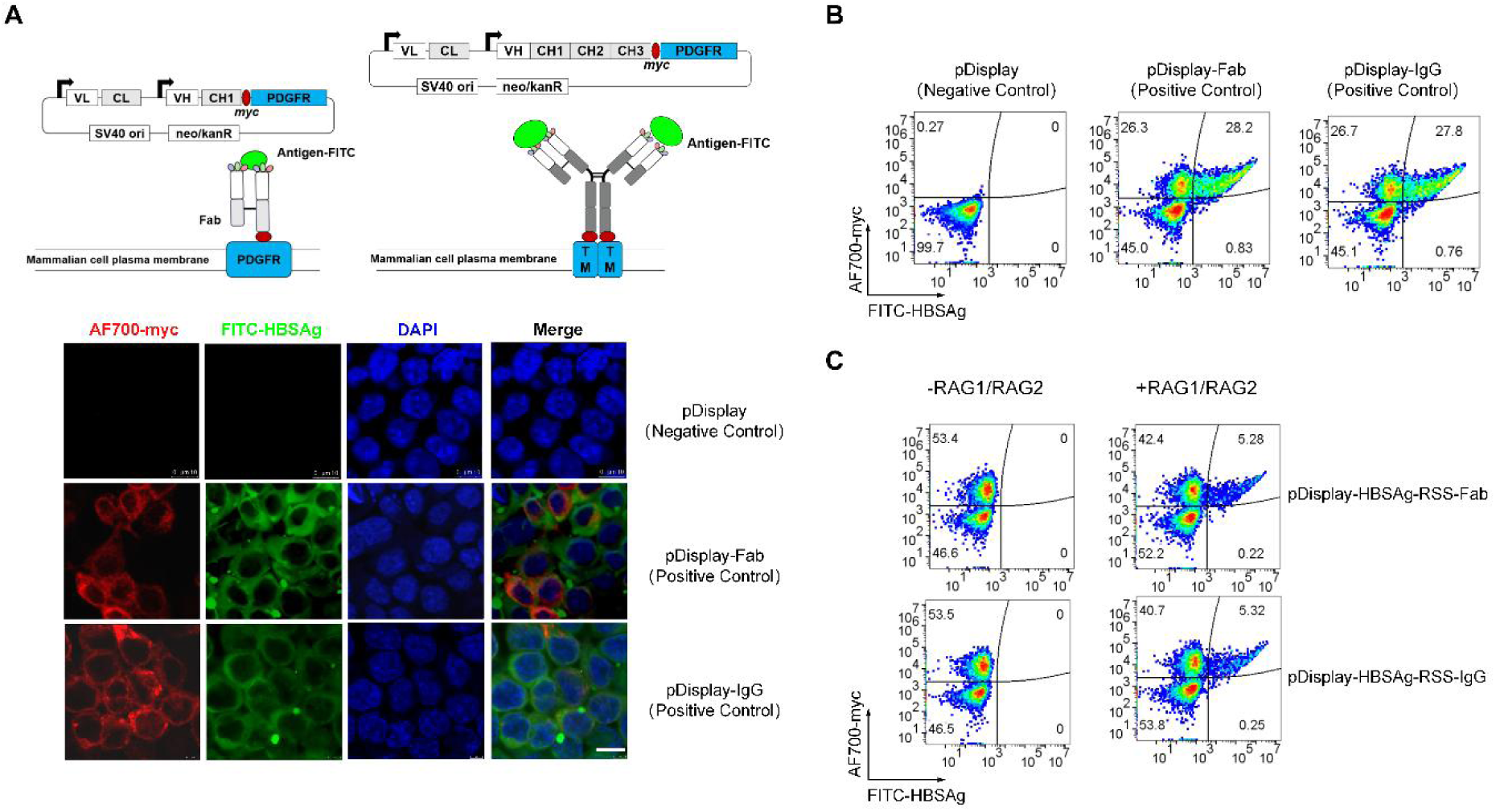
Mammalian cell surface display and flow cytometric analysis of Fab and IgG antibodies. (A) Immuno-fluorescence analysis of antibody surface display and antigen binding. Immunofluorescence staining of cells demonstrating the surface display and functionality of engineered antibodies. Panels show: a negative control transfected with empty pDisplay plasmid (top), successful display and HBSAg-binding of Fab antibodies (middle) and full-length IgG antibodies (bottom). Red fluorescence indicates the myc tag labeled with an AF700 conjugated antibody, green fluorescence shows FITC-conjugated HBSAg antigen, and blue represents DAPI-stained nuclei. The co-localization of red and green signals on the cell membrane confirms successful antibody display and specific antigen-binding capability. Scale bar:10μm. (B) Flow cytometric analysis of surface display efficiency. Pseudocolor dot plots depict the surface expression of the myc-tagged antibodies in cells transfected with empty pDisplay plasmid (left), Fab- displaying construct (middle), or IgG- displaying construct (right). (C) ARESEC-mediated reprogramming yields functional light chains that successfully assemble into antigen-binding Fab (top) and IgG (bottom) antibody formats, both of which demonstrate specific recognition of HBSAg.

**Figure S5.**
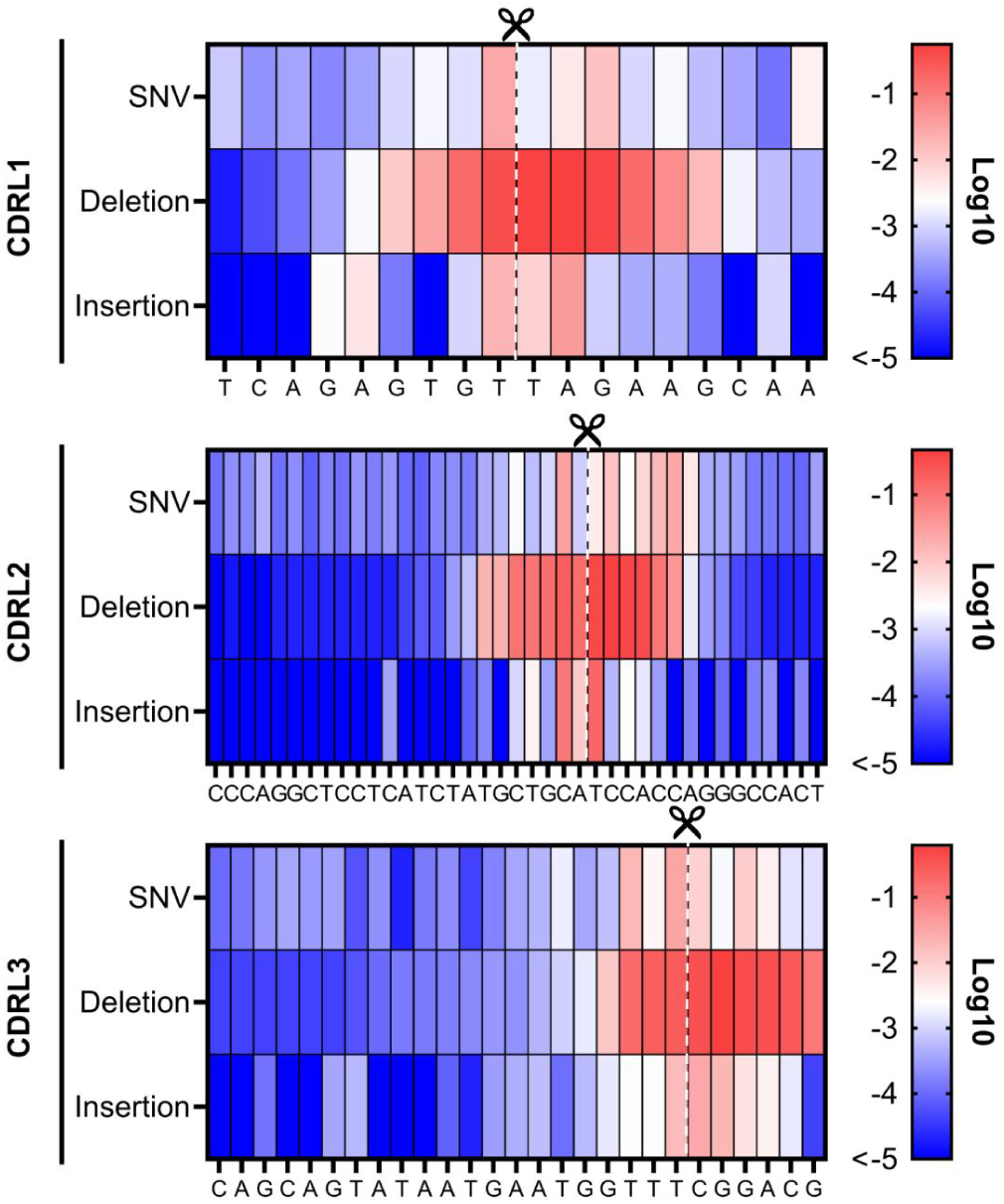
Nucleotide Diversification Profile of CDRL 1/2/3 by High-Throughput Sequencing. High-throughput sequencing analysis of nucleotide-level diversification across CDRL1, CDRL2, and CDRL3 of HBSAg antibody. The heatmap shows log10-transformed values of mutation frequency. Scissors and dashed lines indicate RAG1/2 recognition and cleavage sites.

**Figure S6.**
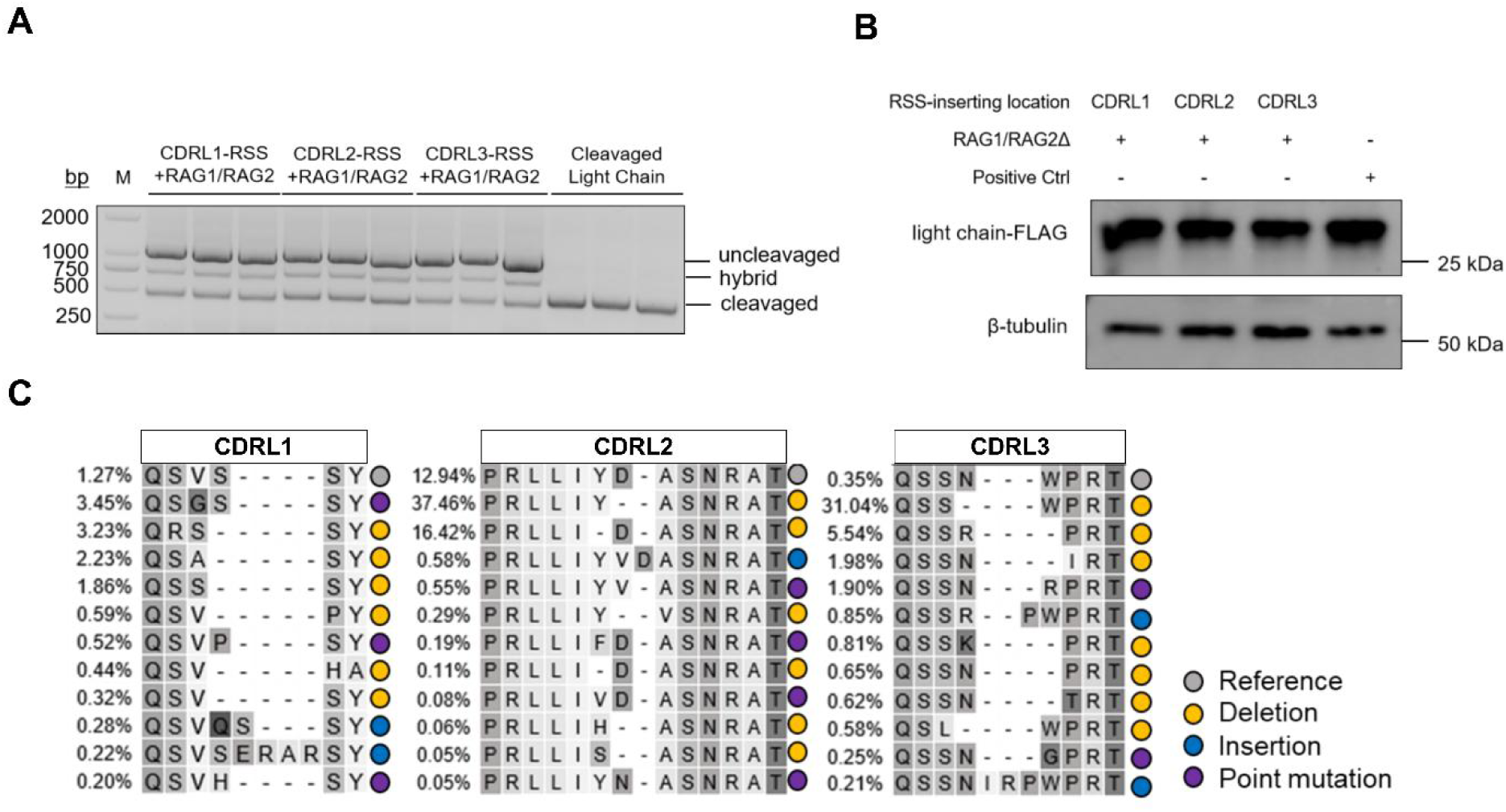
Reprogramming of Nivolumab by the ARESEC Platform, related to Figure 5. (A) PCR analysis of the rearrangement efficiency in CDRL1, CDRL2, and CDRL3 regions. (B) Western blot analysis confirming light chain protein expression after reprogramming. (C) Top 12 most abundant amino acid sequences and their mutation types in Nivolumab following rearrangement by ARESEC. The percentage to the left of each sequence indicates the proportion of reads in the DNA next-generation sequencing data. The sequences are color-coded: gray for the reference sequence, yellow for deletions, blue for insertions, and purple for amino acid point mutations.

**Figure S7.**
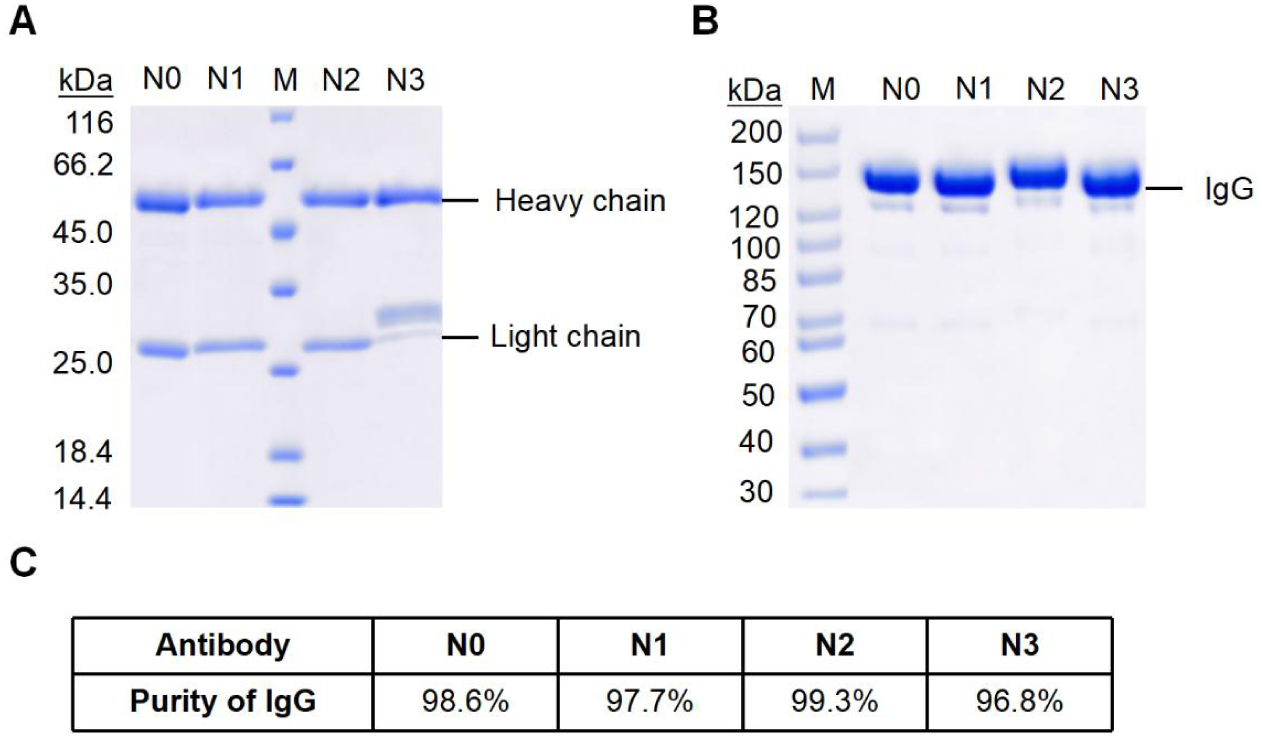
Analysis of integrity, purity, and aggregation state of Nivolumab and its reprogrammed variants after purification, related to Figure 5. (A) SDS-PAGE analysis under reducing conditions. Lane M: Protein molecular weight marker; Lanes N0-N3: Purified antibody samples, showing bands at approximately 50 kDa (Heavy Chain) and 25 kDa (Light Chain). (B) SDS-PAGE analysis under non-reducing conditions. Lane M: Protein molecular weight marker; Lanes N0-N3: Purified antibody samples, showing a single major band at approximately 150 kDa, corresponding to the intact IgG molecule. (C) Purity of IgG, analyzed by Size-exclusion chromatography (SEC-HPLC).

**Figure S8.**
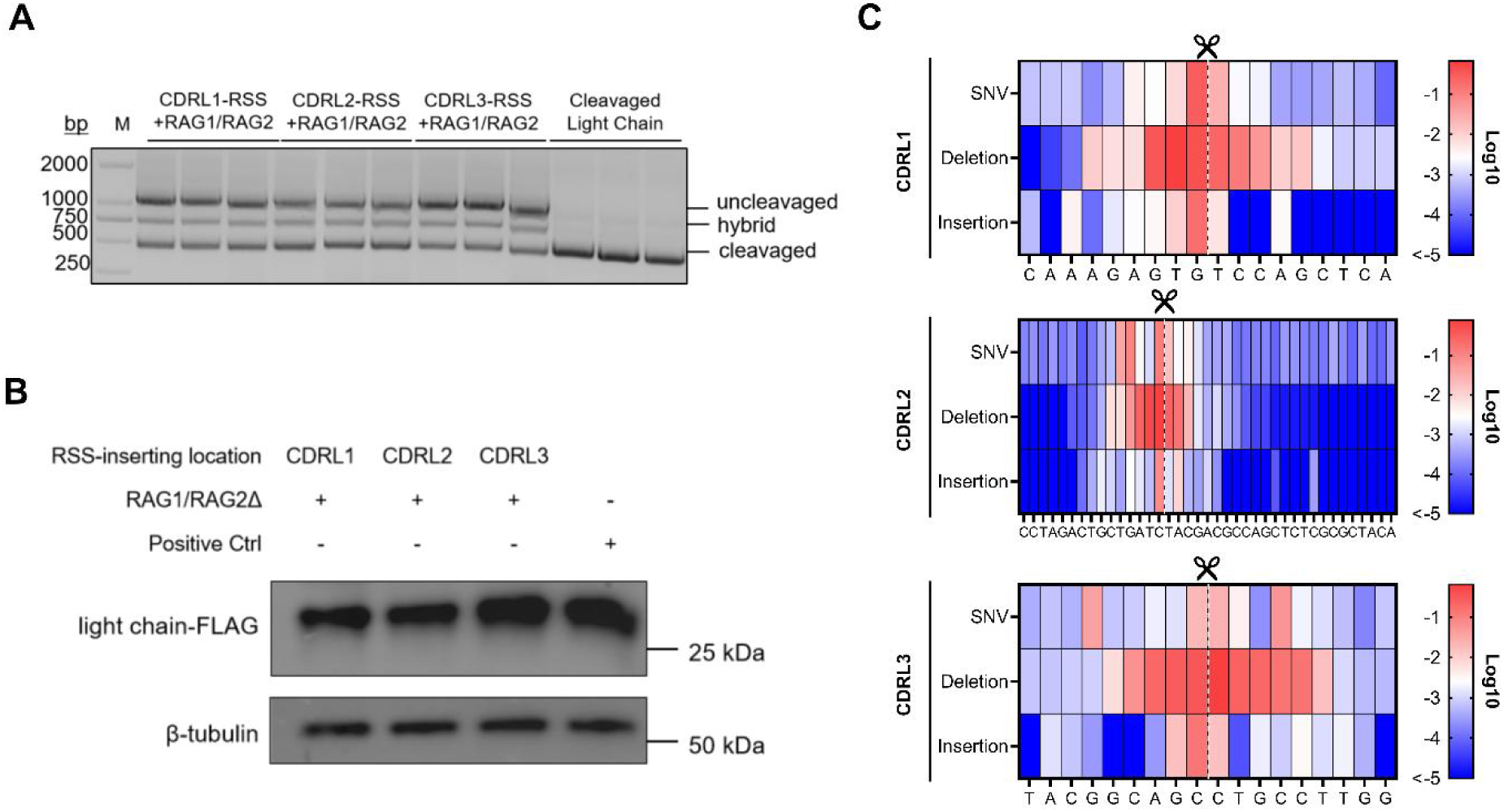
Reprogramming of Durvalumab by the ARESEC Platform, related to Figure 6. (A) PCR analysis of the rearrangement efficiency in CDRL1, CDRL2, and CDRL3 regions. (B) Western blot analysis confirming light chain protein expression after reprogramming. (C) Sequencing analysis of the distribution and frequency of nucleotide mutation types at each position in CDRL1, CDRL2, and CDRL3 following Durvalumab rearrangement by ARESEC. The heatmap shows log₁₀-transformed values of mutation frequency. Scissors and dashed lines indicate RAG1/2 recognition and cleavage sites.

## References

[1] Y. H. Zhang, T. C. Cheng, G. R. Huang, et al.,“Transposon molecular domestication and the evolution of the RAG recombinase”, Nature (2019), 569 (7754), 79, 10.1038/s41586-019-1093-7.

[2] G. W. Litman, J. P. Rast, S. D. Fugmann,“The origins of vertebrate adaptive immunity”, Nature Reviews Immunology (2010), 10 (8), 543, 10.1038/nri2807.

[3] C. Liu, Y. H. Zhang, C. C. Liu, et al.,“Structural insights into the evolution of the RAG recombinase”, Nature Reviews Immunology (2022), 22 (6), 353, 10.1038/s41577-021-00628-6.

[4] B. Bertocci, A. De Smet, J. C. Weill, et al.,“Nonoverlapping functions of DNA polymerases Mu, lambda, and terminal deoxynucleotidyltransferase during immunoglobulin V(D)J recombination in vivo”, Immunity (2006), 25 (1), 31, 10.1016/j.immuni.2006.04.013.

[5] H. Lu, K. Schwarz, M. R. Lieber,“Extent to which hairpin opening by the Artemis: DNA-PKcs complex can contribute to junctional diversity in V(D)J recombination”, Nucleic Acids Research (2007), 35 (20), 6917, 10.1093/nar/gkm823.

[6] H. B. Chi, M. Pepper, P. G. Thomas,“Principles and therapeutic applications of adaptive immunity”, Cell (2024), 187 (9), 2052, 10.1016/j.cell.2024.03.037.

[7] M. S. Kim, M. Lapkouski, W. Yang, et al.,“Crystal structure of the V(D)J recombinase RAG1-RAG2”, Nature (2015), 518 (7540), 507, 10.1038/nature14174.

[8] H. Lu, N. Shimazaki, P. Raval, et al.,“A biochemically defined system for coding joint formation in V(D)J recombination”, Molecular Cell (2008), 31 (4), 485, 10.1016/j.molcel.2008.05.029.

[9] D. J. Guo, D. K. Dunn-Walters, F. Fraternali, et al.,“ImmunoMatch learns and predicts cognate pairing of heavy and light immunoglobulin chains”, Nature Methods (2025), 10.1038/s41592-025-02913-x.

[10] T. Zuo, A. Gautam, S. Saghaei, et al.,“Somatic hypermutation generates antibody specificities beyond the primary repertoire”, Immunity (2025), 58 (6), 10.1016/j.immuni.2025.04.014.

[11] Y. Y. Wang, S. X. Zhang, X. R. Yang, et al.,“Mesoscale DNA feature in antibody-coding sequence facilitates somatic hypermutation”, Cell (2023), 186 (10), 2193, 10.1016/j.cell.2023.03.030.

[12] O. B. Giorgetti, C. P. O’Meara, M. Schorpp, et al.,“Origin and evolutionary malleability of T cell receptor a diversity”, Nature (2023), 619 (7968), 193, 10.1038/s41586-023-06218-x.

[13] Y. H. Zhang, K. Shetty, M. D. Surleac, et al.,“Mapping and Quantitation of the Interaction between the Recombination Activating Gene Proteins RAG1 and RAG2”, The Journal of Biological Chemistry (2015), 290 (19), 11802, 10.1074/jbc.M115.638627.

[14] M. Lapkouski, W. Chuenchor, M. S. Kim, et al.,“Assembly Pathway and Characterization of the RAG1/2-DNA Paired and Signal-end Complexes”, The Journal of Biological Chemistry (2015), 290 (23), 14618, 10.1074/jbc.M115.641787.

[15] D. C. vanGent, D. A. Ramsden, M. Gellert,“The RAG1 and RAG2 proteins establish the 12/23 rule in V(D)J recombination”, Cell (1996), 85 (1), 107, 10.1016/S0092-8674(00)81086-7.

[16] H. Ru, M. G. Chambers, T. M. Fu, et al.,“Molecular Mechanism of V(D)J Recombination from Synaptic RAG1-RAG2 Complex Structures”, Cell (2015), 163 (5), 1138, 10.1016/j.cell.2015.10.055.

[17] W. Hoolehan, J. C. Harris, J. N. Byrum, et al.,“An updated definition of V(D)J recombination signal sequences revealed by high-throughput recombination assays”, Nucleic Acids Research (2022), 50 (20), 11696, 10.1093/nar/gkac1038.

[18] C. J. Wu, Y. Y. Dong, X. H. Zhao, et al.,“RAG2 involves the Igκ locus demethylation during B cell development”, Molecular Immunology (2017), 88, 125, 10.1016/j.molimm.2017.06.026.

[19] A. G. W. Matthews, A. J. Kuo, S. Ramón-Maiques, et al.,“RAG2 PHD finger couples histone H3 lysine 4 trimethylation with V(D)J recombination”, Nature (2007), 450 (7172), 1106, 10.1038/nature06431.

[20] B. A. Helmink, B. P. Sleckman,“The Response to and Repair of RAG-Mediated DNA Double-Strand Breaks”, Annual Review of Immunology (2012), 30, 175, 10.1146/annurev-immunol-030409-101320.

[21] Y. M. Ma, U. Pannicke, K. Schwarz, et al.,“Hairpin opening and overhang processing by an Artemis/DNA-dependent protein kinase complex in nonhomologous end joining and V(D)J recombination”, Cell (2002), 108 (6), 781, 10.1016/S0092-8674(02)00671-2.

[22] T. C. Deiss, M. Vadnais, F. Wang, et al.,“Immunogenetic factors driving formation of ultralong VH CDR3 in antibodies”, Cellular & Molecular Immunology (2019), 16 (1), 53, 10.1038/cmi.2017.117.

[23] J. L. Xu, M. M. Davis,“Diversity in the CDR3 region of V is sufficient for most antibody specificities”, Immunity (2000), 13 (1), 37, 10.1016/S1074-7613(00)00006-6.

[24] J. A. Douthwaite, S. Sridharan, C. Huntington, et al.,“Affinity maturation of a novel antagonistic human monoclonal antibody with a long V CDR3 targeting the Class A GPCR formyl-peptide receptor 1”, Mabs (2015), 7 (1), 152, 10.4161/19420862.2014.985158.

[25] E. K. Makowski, T. X. Wang, J. M. Zupancic, et al.,“Optimization of therapeutic antibodies for reduced self-association and non-specific binding via interpretable machine learning”, Nature Biomedical Engineering (2024), 8 (1), 10.1038/s41551-023-01074-6.

[26] D. M. Mason, S. Friedensohn, C. R. Weber, et al.,“Optimization of therapeutic antibodies by predicting antigen specificity from antibody sequence via deep learning”, Nature Biomedical Engineering (2021), 5 (6), 600, 10.1038/s41551-021-00699-9.

[27] E. K. Makowski, P. C. Kinnunen, J. Huang, et al.,“Co-optimization of therapeutic antibody affinity and specificity using machine learning models that generalize to novel mutational space”, Nature Communications (2022), 13 (1), 10.1038/s41467-022-31457-3.

[28] C. Y. Yi, X. Y. Sun, J. Ye, et al.,“Key residues of the receptor binding motif in the spike protein of SARS-CoV-2 that interact with ACE2 and neutralizing antibodies”, Cellular & Molecular Immunology (2020), 17 (6), 621, 10.1038/s41423-020-0458-z.

[29] A. C. Hunt, B. Voegeli, A. O. Hassan, et al.,“A rapid cell-free expression and screening platform for antibody discovery”, Nature Communications (2023), 14 (1), 10.1038/s41467-023-38965-w.

[30] E. K. Wagner, K. P. Carter, Y. W. Lim, et al.,“Report High-throughput specificity profiling of antibody libraries using ribosome display and microfluidics”, Cell Reports Methods (2024), 4 (12), 10.1016/j.crmeth.2024.100934.

[31] X. L. Zhuang, Z. Wang, J. S. Fan, et al.,“Structure-guided and phage-assisted evolution of a therapeutic anti-EGFR antibody to reverse acquired resistance”, Nature Communications (2022), 13 (1), 10.1038/s41467-022-32159-6.

[32] B. M. Petersen, M. B. Kirby, K. M. Chrispens, et al.,“An integrated technology for quantitative wide mutational scanning of human antibody Fab libraries”, Nature Communications (2024), 15 (1), 10.1038/s41467-024-48072-z.

[33] Y. Zhang,“Evolution of phage display libraries for therapeutic antibody discovery”, Mabs (2023), 15 (1), 10.1080/19420862.2023.2213793.

[34] H. A. Parray, S. Shukla, S. Samal, et al.,“Hybridoma technology a versatile method for isolation of monoclonal antibodies, its applicability across species, limitations, advancement and future perspectives”, International Immunopharmacology (2020), 85, 10.1016/j.intimp.2020.106639.

[35] H. Satofuka, S. Abe, T. Moriwaki, et al.,“Efficient human-like antibody repertoire and hybridoma production in trans-chromosomic mice carrying megabase-sized human immunoglobulin loci”, Nature Communications (2022), 13 (1), 10.1038/s41467-022-29421-2.

[36] N. Lin, K. Miyamoto, T. Ogawara, et al.,“Epitope binning for multiple antibodies simultaneously using mammalian cell display and DNA sequencing”, Communications Biology (2024), 7 (1), 10.1038/s42003-024-06363-7.

[37] E. Kang, C. Kadoch, J. L. Rubenstein, et al.,“A functional mammalian display screen identifies rare antibodies that stimulate NK cell-mediated cytotoxicity”, Proceedings of the National Academy of Sciences of the United States of America (2021), 118 (31), 10.1073/pnas.2104099118.

[38] J. E. Shin, A. J. Riesselman, A. W. Kollasch, et al.,“Protein design and variant prediction using autoregressive generative models”, Nature Communications (2021), 12 (1), 10.1038/s41467-021-22732-w.

[39] K. Fischer, A. Lulla, T. Y. So, et al.,“Rapid discovery of monoclonal antibodies by microfluidics-enabled FACS of single pathogen-specific antibody-secreting cells”, Nature Biotechnology (2025), 43 (6), 10.1038/s41587-024-02346-5.

[40] J. S. James, J. B. Dai, W. L. Chew, et al.,“The design and engineering of synthetic genomes”, Nature Reviews Genetics (2025), 26 (5), 298, 10.1038/s41576-024-00786-y.

[41] L. M. Mindrebo, H. J. Liu, G. Ozorowski, et al.,“Fully synthetic platform to rapidly generate tetravalent bispecific nanobody-based immunoglobulins”, Proceedings of the National Academy of Sciences of the United States of America (2023), 120 (24), 10.1073/pnas.2216612120.

[42] L. Zhong, Q. Zhang, N. Lu, et al.,“The conjugation-associated linear-BAC iterative assembling (CALBIA) method for cloning 2.1-Mb human chromosomal DNAs in bacteria”, Cell Research (2025), 35 (4), 309, 10.1038/s41422-024-01063-7.

[43] J. Y. Chen, N. Singh, J. X. Lu, et al.,“Artificial intelligence-powered biofoundries for protein engineering and metabolic engineering”, Current Opinion in Biotechnology (2025), 96, 10.1016/j.copbio.2025.103380.

[44] J. H. Ministro, S. S. Oliveira, J. G. Oliveira, et al.,“Synthetic antibody discovery against native antigens by CRISPR/Cas9-library generation and endoplasmic reticulum screening”, Applied Microbiology and Biotechnology (2020), 104 (6), 2501, 10.1007/s00253-020-10423-3.

[45] D. M. Mason, C. R. Weber, C. Parola, et al.,“High-throughput antibody engineering in mammalian cells by CRISPR/Cas9-mediated homology-directed mutagenesis”, Nucleic Acids Research (2018), 46 (14), 7436, 10.1093/nar/gky550.

[46] K. Göpfrich, M. Platten, F. Frischknecht, et al.,“Bottom-up synthetic immunology”, Nature Nanotechnology (2024), 19 (11), 1587, 10.1038/s41565-024-01744-9.

[47] F. Caliendo, M. Dukhinova, V. Siciliano,“Engineered Cell-Based Therapeutics: Synthetic Biology Meets Immunology”, Frontiers in Bioengineering and Biotechnology (2019), 7, 10.3389/fbioe.2019.00043.

[48] D. G. Schatz, Y. H. Ji,“Recombination centres and the orchestration of V(D)J recombination”, Nature Reviews Immunology (2011), 11 (4), 251, 10.1038/nri2941.

[49] J. Byrum, W. A. Rodgers, K. Rodgers,“Full Length RAG2 Expression Enhances the DNA Damage Response in Pre-B Cells”, The Journal of Immunology (2018), 200 (1), 10.1016/j.imbio.2021.152089

[50] L. Zhang, T. L. Reynolds, X. C. Shan, et al.,“Coupling of V(D)J Recombination to the Cell Cycle Suppresses Genomic Instability and Lymphoid Tumorigenesis”, Immunity (2011), 34 (2), 163, 10.1016/j.immuni.2011.02.003.

[51] C. Y. Wang, K. B. Thudium, M. H. Han, et al.,“Characterization of the Anti-PD-1 Antibody Nivolumab, BMS-936558, and Toxicology in Non-Human Primates”, Cancer Immunology Research (2014), 2 (9), 846, 10.1158/2326-6066.Cir-14-0040.

[52] S. G. Tan, H. Zhang, Y. Chai, et al.,“An unexpected N-terminal loop in PD-1 dominates binding by nivolumab”, Nature Communications (2017), 8, 10.1038/ncomms14369.

[53] J. Y. Lee, H. T. Lee, W. Shin, et al.,“Structural basis of checkpoint blockade by monoclonal antibodies in cancer immunotherapy”, Nature Communications (2016), 7, 10.1038/ncomms13354.

[54] A. P. Bejarano, O. M. Pérez,“Successful treatment with durvalumab: A case report and review”, Journal of Cancer Research and Therapeutics (2023), 19 (2), 470, 10.4103/jcrt.jcrt_1430_21.

[55] J. Alvarez-Argote, C. A. Dasanu,“Durvalumab in cancer medicine: a comprehensive review”, Expert Opinion on Biological Therapy (2019), 19 (9), 927, 10.1080/14712598.2019.1635115.

[56] S. M. Christie, C. Fijen, E. Rothenberg,“V(D)J Recombination: Recent Insights in Formation of the Recombinase Complex and Recruitment of DNA Repair Machinery”, Frontiers in Cell and Developmental Biology (2022), 10, 10.3389/fcell.2022.886718.

[57] A. P. Cazier, J. Son, S. Yellayi, et al.,“Generating combinatorial diversity via engineered V(D)J-like recombination in *Saccharomyces cerevisiae*”, Nature Communications (2025), 16 (1), 10.1038/s41467-025-61206-1.

[58] J. A. E. Repasky, E. Corbett, C. Boboila, et al.,“Mutational analysis of terminal deoxynucleotidyltransferase-mediated N-nucleotide addition in V(D)J recombination”, J Immunol (2004), 172 (9), 5478, 10.4049/jimmunol.172.9.5478.

[59] L. Liu, J. Li, M. Cisneros-Aguirre, et al.,“Dynamic assemblies and coordinated reactions of non-homologous end joining”, Nature (2025), 643 (8072), 10.1038/s41586-025-09078-9.

[60] S. Ding, X. H. Wu, G. Li, et al.,“Efficient transposition of the piggyBac(PB) transposon in mammalian cells and mice”, Cell (2005), 122 (3), 473, 10.1016/j.cell.2005.07.013.

[61] S. Yamamoto, S. Matsumoto, A. Goto, et al.,“Quantitative PCR methodology with a volume-based unit for the sophisticated cellular kinetic evaluation of chimeric antigen receptor T cells”, Scientific Reports (2020), 10 (1), 10.1038/s41598-020-74927-8.

[62] T. T. Gan, Y. H. Wang, Y. Liu, et al.,“RAG2 abolishes RAG1 aggregation to facilitate V(D)J recombination”, Cell Reports (2021), 37 (2), 10.1016/j.celrep.2021.109824.

[63] T. H. Bruun, V. Grassmann, B. Zimmer, et al.,“Mammalian cell surface display for monoclonal antibody-based FACS selection of viral envelope proteins”, Mabs (2017), 9 (7), 1052, 10.1080/19420862.2017.1364824.

[64] C. Zhou, F. W. Jacobsen, L. Cai, et al.,“Development of a novel mammalian cell surface antibody display platform”, Mabs (2010), 2 (5), 508, 10.4161/mabs.2.5.12970.

[65] R. R. Beerli, M. Bauer, R. B. Buser, et al.,“Isolation of human monoclonal antibodies by mammalian cell display”, Proceedings of the National Academy of Sciences of the United States of America (2008), 105 (38), 14336, 10.1073/pnas.0805942105.

[67] M. Tabasi, S. Eybpoosh, S. Bouzari,“Development of an indirect ELISA based on whole cell lysates for detection of IgM anti-antibodies in human serum”, *Comparative Immunology*, Microbiology and Infectious Diseases (2019), 63, 87, 10.1016/j.cimid.2019.01.007.

[68] C. Dacon, C. Tucker, L. H. Peng, et al.,“Broadly neutralizing antibodies target the coronavirus fusion peptide”, Science (2022), 377 (6607), 728, 10.1126/science.abq3773.

[69] T. L. Kirley, A. B. Norman,“Unfolding of IgG domains detected by non-reducing SDS-PAGE”, Biochemical and Biophysical Research Communications (2018), 503 (2), 944, 10.1016/j.bbrc.2018.06.100.

[70] Y. P. Feng, M. Yuan, J. M. Powers, et al.,“Broadly neutralizing antibodies against sarbecoviruses generated by immunization of macaques with an AS03-adjuvanted COVID-19 vaccine”, Science Translational Medicine (2023), 15 (695), 10.1126/scitranslmed.adg7404.

